# Hierarchical Lineage Tracing Reveals Diverse Pathways of AML Treatment Resistance

**DOI:** 10.1101/2025.02.27.640600

**Authors:** Rachel Saxe, Hannah Stuart, Abigail Marshall, Fahiima Abdullahi, Zoë Chen, Francesco Emiliani, Aaron McKenna

## Abstract

Cancer cells adapt to treatment, leading to the emergence of clones that are more aggressive and resistant to anti-cancer therapies. We have a limited understanding of the development of treatment resistance as we lack technologies to map the evolution of cancer under the selective pressure of treatment. To address this, we developed a hierarchical, dynamic lineage tracing method called FLARE (Following Lineage Adaptation and Resistance Evolution). We use this technique to track the progression of acute myeloid leukemia (AML) cell lines through exposure to Cytarabine (AraC), a front-line treatment in AML, in vitro and in vivo. We map distinct cellular lineages in murine and human AML cell lines predisposed to AraC persistence and/or resistance via the upregulation of cell adhesion and motility pathways. Additionally, we highlight the heritable expression of immunoproteasome 11S regulatory cap subunits as a potential mechanism aiding AML cell survival, proliferation, and immune escape in vivo. Finally, we validate the clinical relevance of these signatures in the TARGET-AML cohort, with a bisected response in blood and bone marrow. Our findings reveal a broad spectrum of resistance signatures attributed to significant cell transcriptional changes. To our knowledge, this is the first application of dynamic lineage tracing to unravel treatment response and resistance in cancer, and we expect FLARE to be a valuable tool in dissecting the evolution of resistance in a wide range of tumor types.

## Introduction

Acute myeloid leukemia (AML) is a heterogeneous hematological malignancy that has served as a model of clonal evolution in response to treatment^1–4^. AML subtypes and risk categories have long been primarily defined by cytogenetics, mutation status, and structural genomic variations. However, with recent advances in RNA sequencing, transcriptome profiling has begun to uncover further leukemic disease states, dependencies, and prognostic signatures^5–7^. Upon diagnosis, most patients are treated with induction therapy, including an aggressive cycle of Cytarabine (AraC). While ∼80% of patients reach remission following induction, most will relapse^8^. Single-cell RNA sequencing (scRNA-seq) of matched pre– and post-treatment patient bone marrow samples show patients have dramatic disease landscape changes following AraC treatment^2^.

It remains unclear how resistance to AraC evolves in many AML patients. Are AraC-resistant subclones pre-existing, or does resistance emerge de novo in rare subpopulations? Does the resistant phenotype pass through multiple intermediate states or emerge in a punctuated evolution? Previous studies have investigated the origin of chemoresistant cell populations in various cancer types^9–12^. Shaffer et al. concluded that while rare subpopulations of cells exhibit pre-resistant-like phenotypes, melanoma cells could transiently shift to a resistant phenotype upon treatment in vitro^11^. In contrast, Yan et al. identified a subpopulation of TP53-mutated AML cells in treatment-naïve samples believed to be the source of future AraC-resistant populations^9^. These studies highlight that treatment failure and resistance are likely driven by diverse disease, drug, and likely, patient-specific mechanisms. While individual patients may display unique lineage developments and gene expression profiles, pathway alterations that arise frequently can be exploited for treatment response prediction, patient stratification, and development of targeted therapies.

We set out to develop a cellular lineage tracing system to record the evolutionary steps of AraC treatment resistance in AML. Here, we present FLARE (Following Lineage Adaptation and Resistance Evolution), a multi-tiered lineage tracing approach, which we use to identify clonal patterns of drug response and expansion in vitro and in vivo. Using advances in CRISPR lineage tracing, we track the evolution of cell fate independently of cell state, recording the relationship between cells in small, integrated recorders, which can be used to recover cellular ‘phylogenetic’ trees^13,14^. This lineage information alone can uncover treatment response patterns, but when paired with transcriptomic data, it acts as a window into disease evolution. CRISPR lineage recording has been previously used to characterize clonal evolution through tumorigenesis and metastasis, yet the effect of treatment has yet to be explored^14,15^.

We use FLARE to uncover multiple heritable and lineage-specific gene modules associated with drug response with matched outcomes in pediatric AML patient survival. FLARE lineage patterns uncovered a highly heritable gene signature in treatment-naive AML cells involved in cell-adhesion-mediated drug resistance (CAM-DR) pathways, the expression of which leads to lineage branch survival and expansion under AraC exposure. In addition, we identified an AML lineage-dependent upregulation of immunoproteasome subunits, with further upregulation leading to improved expansion in vivo. Finally, we validated the clinical relevance of our findings using RNA sequencing data from the pediatric TARGET AML cohort and discover bone marrow and peripheral blood-specific expression profiles significantly associated with patient outcomes.

## Results

### Hierarchical lineage recording in AML

We set out to develop a dynamic cellular lineage tracing system to model the fitness landscape of AML in vitro and in vivo. Expanding on our previous macsGESTALT lineage tracing system^14^, we created a hierarchical system, capturing lineage information at three levels (**Figure 1A-B**). At the top level, FLARE encodes founder populations using a unique combination of static tags accompanying the lineage recorders when integrated into the genome. Within each labeled founder population, individual clones are divided into sub-populations with a second round of tags integrated with the Cas9 cassette. This Cas9 integration also begins the dynamic lineage recording, creating insertions and deletions (indels) within the lineage recorders, capturing sub-clonal lineage information. We can then use computational methods to generate a phylogenetic tree of cell relationships within these founder and clonal populations from these indels. This three-level structure (founder, clone, and subclonal lineage information) allows FLARE to track dynamics across time and to compare sister populations in both the control and treatment arms, in vitro and in vivo.

**Figure 1:**
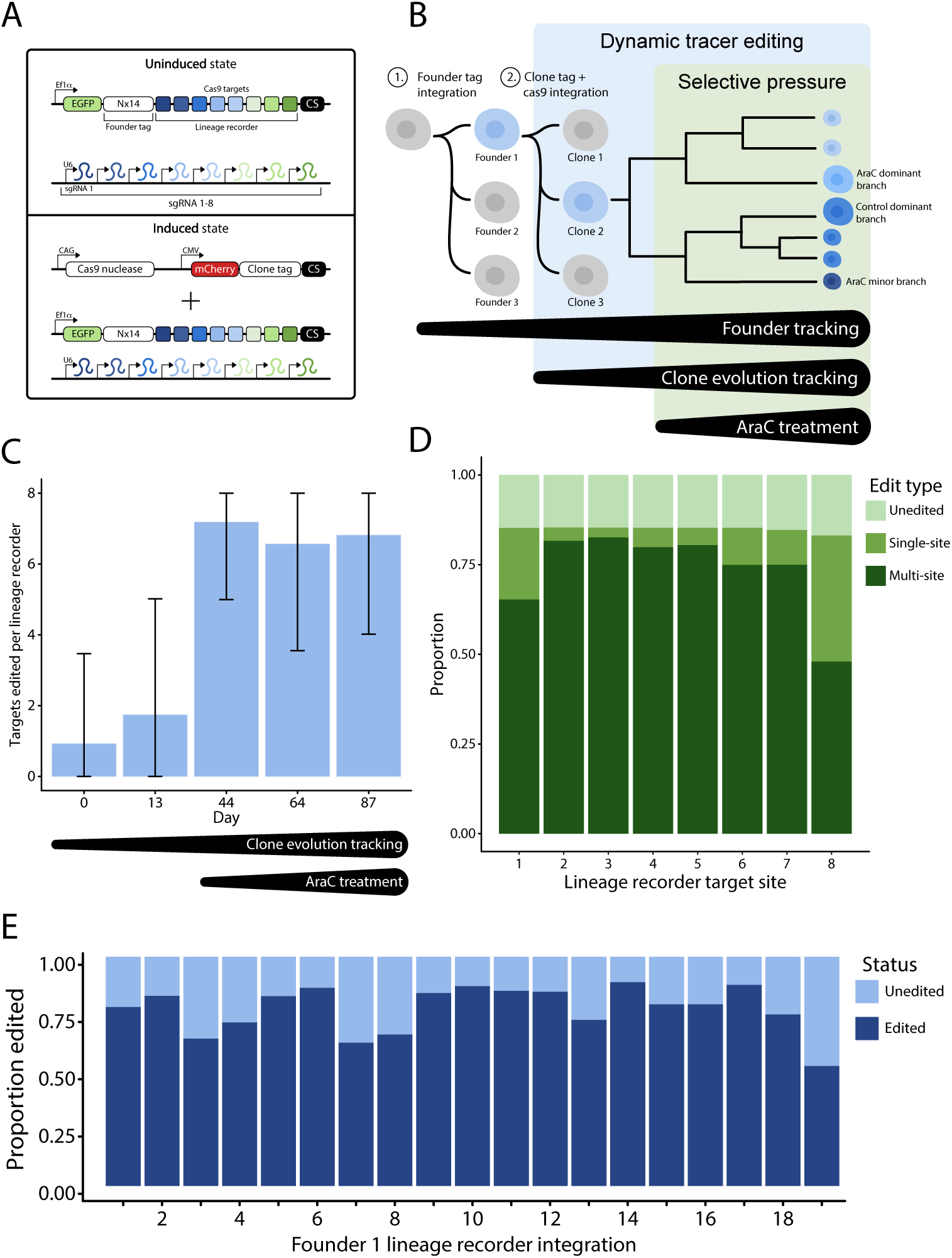
FLARE lineage tracing system allows for hierarchical tracking of cell populations through time. (**A**) Constructs required for FLARE lineage tracing. The top panel includes FLARE-miniV10-10X-UMI (founder tag + lineage recorder) and FLARE-pbV10g-neo (Cas9 guide sequences). We then electroporate the FLARE-pbCas9-tag-mCherry into cells (bottom panel), tagging each clone and activating CRISPR lineage recording. (**B**) The resulting cells record lineage information at 3 levels, (1) founder, (2) clonal, and (3) subclonal (CRISPR recordings). AraC treatment selective pressure is introduced following the initiation of lineage recorder editing. (**C**) Lineage recorders are readily edited following the introduction of Cas9. Editing rates plateau after around 44 days of Cas9 exposure. (**D**) Following 87 days of Cas9 exposure lineage recorder edits are primarily composed of multi-site editing events, with a smaller proportion of edits impacting single sites. Around 15% of lineage recorder sites remain unedited through 87 days of editing. (**E**) Editing rates are consistent across lineage recorder integrations within the C1498-FLARE Founder 1 population.

We first engineered this approach in the C1498 cell line, a syngeneic mouse model of AML compatible with transplantation into immune-competent with transplantation into immune-competent C57BL/6 mice. Despite being long-passaged, C1498 cells are known to be heterogeneous, making them an optimal model for investigating the diversity of clonal treatment response^16^. We established 11 founder populations from individually isolated cells through successive engineering rounds (**Figure S1A-B)**. Sequencing revealed a unique set of founder tags for each population, varying from 19 to 30 integrations (**Figure S1C**). These isolated founders are heterogeneous, with a wide range of growth kinetics (**Figure S1D**). We found no putative driver mutation differences between individual founders using whole-genome sequencing that would explain this heterogeneity.

### Characterizing lineage recording

Next, we set out to test the lineage recording capacity of our engineered cells. We activated CRISPR recording by integrating the Cas9 construct into the genome (**Figure 1A-B**). After electroporation and sorting, we sampled Cas9-positive C1498-FLARE cell populations at days 13, 44, 64, and 87 and sequenced the lineage recorders to characterize Cas9 editing (**Figure 1C**). Recording proportions suggest the bulk of lineage recorder editing occurs between day 13 (22.61% of recorders contain >= 1 edit) and 44 (99.97% >= 1 edit). Editing rates remained high through at least day 87 (85.93% >= 1 edit), with continuing fluctuations in editing patterns due to population drift. Editing patterns were comparable between lineage recorder target sites (**Figure 1D)**, and most edits span multiple target sites with a smaller proportion of edits affecting individual targets. Approximately 15% of sequenced lineage recorder targets remained unedited, indicating our ability to trace lineage over at least 87 days without saturation.

We further characterized the response of individual recorder integrations with a single founder group, founder 1 (F1), containing 19 unique integrations. All integrations were active recorders displaying diverse editing outcomes (**Figure 1E**). For all founder populations assessed, individual recording events tended to be large, spanning 3.075 targets on average, and each recorder contained an average of 2.57 individual recording events. While edits consisting of large deletions can lead to loss of lineage information, we could still reliably reconstruct lineage trees with an average depth of 5.566 nodes over relevant timescales. We attribute this to this system’s high lineage recording capacity, with engineered cells containing between 152 and 240 CRISPR target sites per cell.

### Clonal treatment response and tree generation in vitro and in vivo

Confident in our ability to record and recover cellular lineage information, we set out to characterize C1498-FLARE population dynamics in response to the chemotherapy AraC in vitro and in vivo (**Figure 2A)**. To recreate the heterogeneity observed in leukemia patients, we pooled four C1498 founder groups for our starting population, which was then used for both in vitro and in vivo AraC treatment.

**Figure 2:**
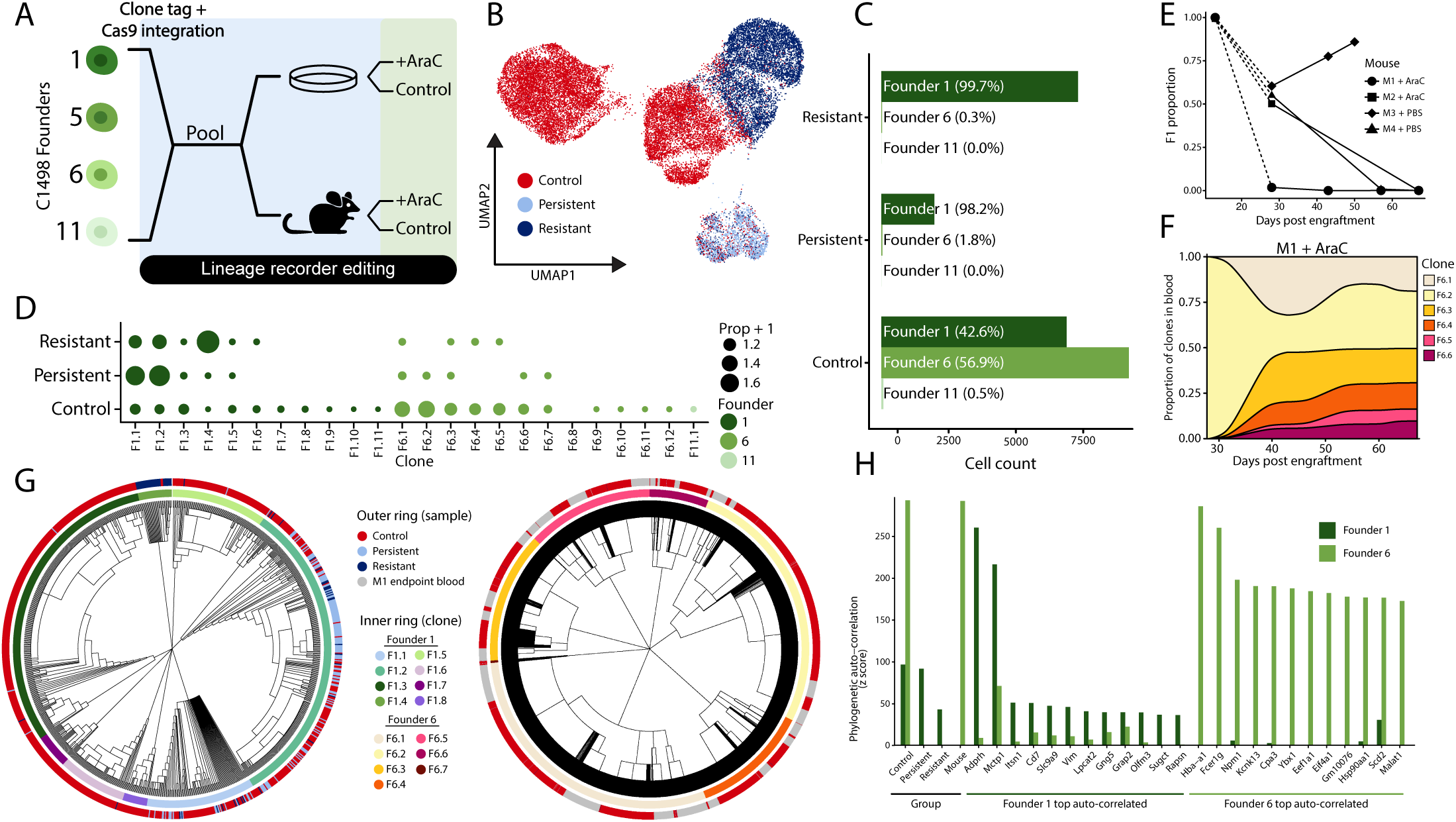
FLARE lineage tracing tracks founder and clone-level evolution in response to AraC treatment in vitro and in vivo. (**A**) Experimental design of lineage tracing C1498-FLARE AML cells in vitro and in vivo. (**B**) Single-cell sequencing of C1498-FLARE cells following 43 days of in vitro AraC treatment. Cells cluster into distinct groups associated with their sample (red-no AraC; dark blue – AraC resistant flask; light blue – AraC persistent flask). (**C**) Founder proportions in each sample display increased survival of F1 cells under AraC treatment in both the resistant and persistent populations. (**D**) Clone populations detected within each sample, ordered by founder and clone size. (**E**) Founder and clone tags detected in blood draws from mice engrafted with C1498-FLARE cells were used to calculate the proportion of circulating leukemic cells belonging to F1 compared to F6. Day 13 proportions were computed using founder tag sequencing and are connected to day 28 with dashed lines. All other timepoint proportions were calculated using clone tag sequencing, normalized for differences in the number of integrations. The F1 proportion was defined as the F1 average tag count divided by the sum of F1 and F6 average tag counts. Untreated mouse M3 was euthanized on day 50 due to disease progression. The remaining mice were euthanized on day 67. (**F**) Bulk sequencing of clone tags detected in blood draws from mouse M1 following AraC treatment on day 13. Clone tag reads were used to determine the clonal proportions of leukemic burden in the blood from day 28 to day 67. (**G**) Phylogenetic reconstruction of founders F1 (left) and F6 (right) generated from single-cell sequencing of in vitro C1498-FLARE lineage tracing and single-cell sequencing of endpoint blood and bone marrow from treated mouse M1. (**H**) Heritability of both sample features and genes for founder 1 and founder 6 computed with PATH.

By day 43 of in vitro AraC treatment we observed two distinct growth phenotypes. Our AraC-resistant flask displayed rapid growth and a 10-fold higher ED50 than control cell populations (**Figure S2A**). We also observed a separate flask phenotype, where cells appeared to be “persistent”, displaying minimal proliferation and cell numbers too low for ED50 determination. We sequenced 27,237 single cells from the untreated, resistant, and persistent flasks. Treated cells clustered into distinct groups predominantly separated by flask phenotype (**Figure 2B**). We successfully captured founder tags and lineage recorders from 100% of single cells and clone tags from 31.9%. After filtering, quality control, and lineage assignment, 25,489 single cells were assigned to founder groups, of which 5,468 were successfully assigned to clones. Founder tag sequencing revealed in vitro fitness differences: in untreated flasks, founder 1 (F1) and founder 6 (F6) dominated (43% and 57% of cells, respectively), and few founder 11 (F11) cells were detected (<0.5% of cells). F1 and F6 control cells clustered separately, suggesting distinct founder-specific transcriptomes (**Figure S2B)**. In contrast, treated populations were primarily composed of F1 with few F6 (<0.7%) and zero F11 cells. F5 was outcompeted in all conditions (**Figure 2C)**.

We performed differential gene expression analysis between founder groups, focusing on F1 and F6 in the control setting. F1 lineages were characterized by high expression of Nkain3, Wnt6, and Chl1 (**Table S1**). In contrast, F6 lineages were characterized by high expression of Sox2ot, Tnni1, and Tnnt1 (**Table S2**). GSEA results indicated enrichment of multiple Hallmark pathways in F6 compared to F1, including MYC targets V1, oxidative phosphorylation, and MTORC1 signaling (**Figure S2C**). Enriching these oncogenic pathways likely contributed to the success of F6 in the absence of treatment. Since these pathways have well-established roles in chemoresistance, we expected F6 to display a survival advantage under AraC treatment^17–20^. Unexpectedly, F6 was almost fully out-competed by F1 in the presence of AraC (**Figure 2C)**.

We then explored founder F1’s competitive advantage over F6 and the other founders during treatment. Within F1 populations, there was a distinct and divergent selection of specific clones in AraC-treated and control flasks (**Figure 2D**). The highest proportion of persistent cells belonged to clones F1.1 and F1.2, while the highest proportion of resistant cells belonged to F1.4. F1 cells have a clear survival advantage through in vitro AraC treatment, but the outgrowth of different clones in treated replicates suggests clone-specific responses that contribute to population growth dynamics.

To identify F1 clone-specific treatment responses, we analyzed differentially expressed genes between control, resistant, and persistent-dominant clones (**Table S3**). Our dominant clone in the resistant population, F1.4 (**Figure 2D**), expressed high levels of IgII1 and Evac1 compared to other resistant cells. Clones F1.1 and F1.2 were successful in both persistent and resistant populations; when we compared their control gene expression to all other F1 clones, a transcriptional signature containing Grap2, Trat1, and Ldlrad4 emerged (**Figure S2D**). Trat1 is involved in the regulation of T cell receptors and expression is associated with immune cell infiltration and improved survival in breast, ovarian, and lung cancer ^21–23^, Ldlrad4 is a negative regulator of TGF-beta signaling ^24,25^, and expression promotes the proliferation and migration of hepatic cancer cells and is downregulated in CD34+ cells from patients with MDS ^26^. Consistent with the upregulation of Ldlrad4 expression in F1.1 and F1.2 control cells, sister cells of both clones in the persistent population display high Ldlrad4. We also identified clones with pre-existing lineage-specific expression patterns. Cells belonging to clone F1.3 tended to colocalize and shared high expression of Adprh and Mctp1 (**Figure S2E-F)**. These genes highlighted by differential gene expression analysis do not have well-established roles in AML.

In parallel to our in vitro treatment assay, we injected immune-competent C57BL/6 KH2^27^ mice with 25k of the pooled founder starting population to induce leukemia with lineage tracing capacity (**Figure 2A)**. Mice were then treated with AraC to a dose comparable to that given to patients in one round of 7+3 induction chemotherapy^9^. We used bulk lineage sequencing to estimate founder and clone dynamics from blood draws of circulating leukemia cells. Contrary to our in vitro results, F6 outcompeted other founder populations in vivo in all but one mouse (**Figure 2E**). This bottlenecking occurred sometime between day 28, when all founders were detectable in control and treated mice (**Figure S3A),** and euthanasia on day 67. A selective sweep was nearly complete in treated mouse M1 (>99.9% F6 by day 43), treated mouse M2 (100% F6 by day 67), and untreated mouse M4 (>99.9% F6 by day 57). Control mouse M3’s leukemic burden was composed of >77.5% F1 by day 43, and this increased to >85.9% on day 50 when M3 was euthanized due to symptomatic disease.

We then focused on the within-founder clone tag sequences seen in blood draws. Interestingly, M1 displayed sustained clonal diversity split amongst F6.1, F6.2, F6.3, F6.4, and F6.5, indicating these clones survived AraC treatment and had comparable fitness during post-treatment outgrowth. In contrast, blood samples from M3 were primarily dominated by F1’s clone F1.2 by day 43 (**Figure S3C)**. Blood samples from M2 and M4 did not contain enough overlap with our known clone tags to infer population dynamics, likely representing diversity not captured in our single-cell sequencing at end-point.

To further examine the lineage identities and transcriptomes of cells surviving in vivo treatment, we sequenced 10,473 and 6,344 cells from blood and bone marrow, respectively, of treated mouse M1 displaying sustained F6 clonal diversity. Non-leukemic cells clustered by cell type, and leukemic cells were identified by founder and clone tags (**Figure S3D-E)**. Of the leukemic cells, 6 and 6,053 belonged to founders F1 and F6, respectively. Clone groups were assigned to 2,002 single cells, 702 belonging to clone F6.1. The remaining clone assignments were split between F6.2, F6.3, F6.4, and F6.5 (**Figures 2F, S3B**). We also detected low frequencies of cells belonging to F6.6 and F6.8. These single-cell sequencing results matched the population detected in bulk sequencing, confirming our ability to reconstruct founder and clone patterns from limited samples, such as blood draws, using FLARE lineage tracing.

We then created phylogenetic trees for F1 and F6 using FLARE lineage recorder indels captured with our in vitro and in vivo scRNA-seq (**Figure 2G)**. To identify heritable gene expression patterns associated with the development of AraC resistance, we conducted lineage autocorrelation analysis on founder trees using PATH (**Figure 2H**)^28^. PATH highlighted genes previously identified by differential gene expression analysis as highly cross-correlated with resistant and/or persistent cell identities in the F1 tree (**Figure S3F**). These genes include Olfm3, a gene associated with anoikis resistance in AML cells^29^, and Cd7, the expression of which in AML is a prognostic risk factor^30–32^; both genes were significantly upregulated in AraC persistent cells. Genes such as Adprh and Mctp1, associated with the F1.3 clone branch, were also strongly heritable but expressed primarily in a subset of control cells.

These initial experiments with C1498-FLARE cells allowed us to optimize the lineage tracing system and test our ability to track leukemic lineage in immune-competent mice. With FLARE, we successfully identified highly heritable genes selected for through AraC treatment that contribute to drug persistence and resistance.

### Clonal treatment response and tree generation in human AML cells

We next engineered FLARE lineage recording human HL60 cells, a model of FAB-M2 AML^33^. We seeded 500 HL60-FLARE founder cells, expanded, and split into parallel cultures (**Figure 3A**). This starting pool expands on the transcriptional landscape sampled in C1498-FLARE cells while maintaining sister founder populations in the control and treatment arms. Similarly to C1498-FLARE cells, our resulting HL60-FLARE cells clustered in distinct groups based on AraC treatment in vitro (**Figure 3B**). However, unlike C1498-FLARE cells, AraC-treated HL60-FLARE cells in different flasks clustered together despite different regrowth rates (**Figure S4A**). AraC-treated flask A reached cell concentrations comparable to untreated HL60-FLARE cells by day 80 of AraC treatment, so we posited these cells would align transcriptionally with C1498-FLARE resistant cells. Surprisingly, differential expression analysis uncovered the upregulation of genes shared with AraC-persistent C1498 F1 cells such as VIM, PSD3, CRIP1, and S100A4 (**Table S4**). AraC-treated flask B displayed slower growth dynamics but shared flask A’s transcriptional profile (**Table S5**).

**Figure 3:**
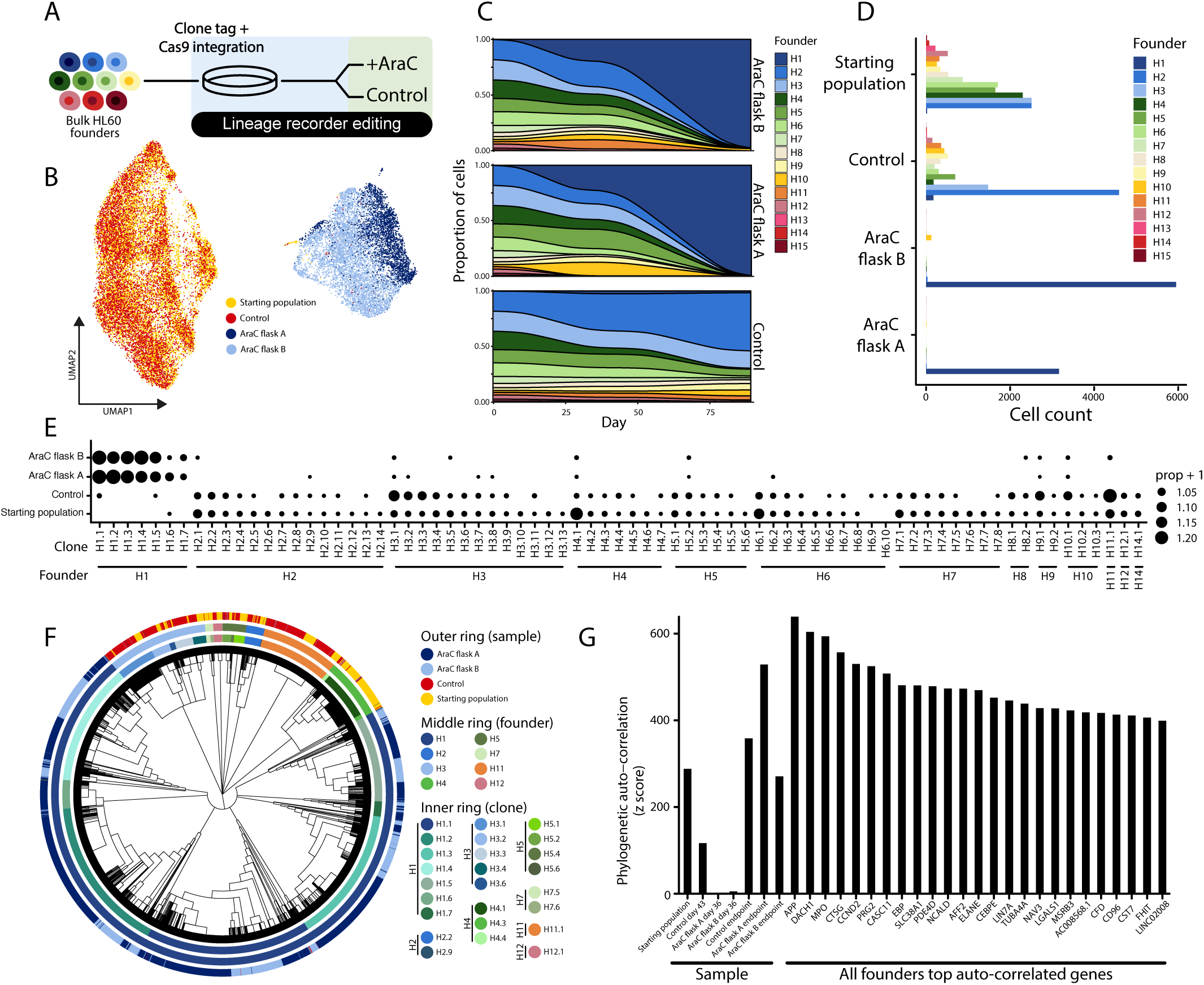
Founder and clone-level selection of founder H1 in response to AraC treatment in human HL60 AML cells. (**A**) Experimental design of HL60-FLARE AML lineage tracing in vitro. (**B**) Single cell sequencing of HL60-FLARE cells following 89 days of in vitro AraC treatment. Cells cluster into distinct groups associated with their sample (yellow – starting population; red – no AraC; dark blue – AraC flask A; light blue – AraC flask B). (**C**) Clone population fluctuations for each flask throughout the experiment. (**D**) Founder proportions in each sample display increased survival of founder H1 cells under AraC treatment in both flask A and B. (**E**) Clone populations for each sample at experimental endpoint, ordered by founder and clone size. (**F**) Phylogenetic reconstruction of HL60-FLARE cells generated from single cell sequencing of in vitro lineage tracing. (**G**) Heritability of both sample features and genes for all HL60-FLARE founders computed via PATH.

Single-cell sequencing of founder and clone tags was used to cluster cells into 15 founder populations (H1-H15), individually composed of 1-14 clones. Sequencing from cells collected at the experiment start, midpoint, and endpoint were used to determine founder proportions over time for each flask. All 15 HL60 founders were detected in the experimental starting population and the control flask at endpoint (**Figure 3C-D**). There was an extensive bottleneck in all cultured populations, regardless of treatment. Untreated endpoint cells were most likely to descend from H2, while both AraC-treated flasks consisted primarily of cells belonging to H1. Both treated flasks maintained diversity within H1 on the clonal level, with most cells being evenly split between clones H1.1, H1.2, H1.3, H1.4, and H1.5 (**Figure 3E**).

The success of H1 in both treated flasks suggests a founder-specific propensity for treatment resistance prior to treatment onset. This is also supported by the comparable fitness of H1 clonal populations, indicating the lack of clone-specific resistance evolution. We used differential gene expression analysis to explore H1-specific expression patterns in the presence and absence of treatment to determine whether H1 cells were consistently differentiated from other founders or were primed for reprogramming following the introduction of AraC. Pre-experiment and endpoint control H1 cells shared the upregulation of 104 genes (**Table S6**). Many of these genes were also upregulated in AraC-resistant H1 cells compared to other founders, including MKX, APBA2, LAMP5, CAMK2D, and NAV3.

We turned to phylogenetic reconstruction and heritability analysis to further dissect the H1’s resistance mechanisms. We used lineage recorder indel patterns to reconstruct phylogenetic trees for each clone population. These clonal trees were merged to create founder-level lineage trees, and founder-level trees were merged to create the HL60 population tree (**Figure 3F**). PATH analysis on the population tree revealed many highly heritable genes associated with evolution with and without AraC, including genes previously implicated in AML prognosis and treatment response, such as MPO (**Figure 3G**). This analysis also indicated significant heritability of genes previously identified as upregulated in treatment-naive H1 cells.

### Identification of pro-leukemic transcriptional programs associated with pre-AraC-resistant lineages

To better understand broad transcriptomic changes over AraC-induced clonal evolution, we generated lineage-dependent gene modules for C1498-FLARE F1 and F6 trees, and the HL60-FLARE population tree using Hotspot (**Figure 4A-B, Table S3, Table S7**)^34^. Jaccard indices revealed significant similarity between a number of HL60 and C1498 modules, indicating shared mechanisms across our two experimental lines (**Figure 4C, S4B-C**). Moreover, expression of these modules within the lineage structure showed founder and clone-specific enrichment (**Figure 4D**).

**Figure 4:**
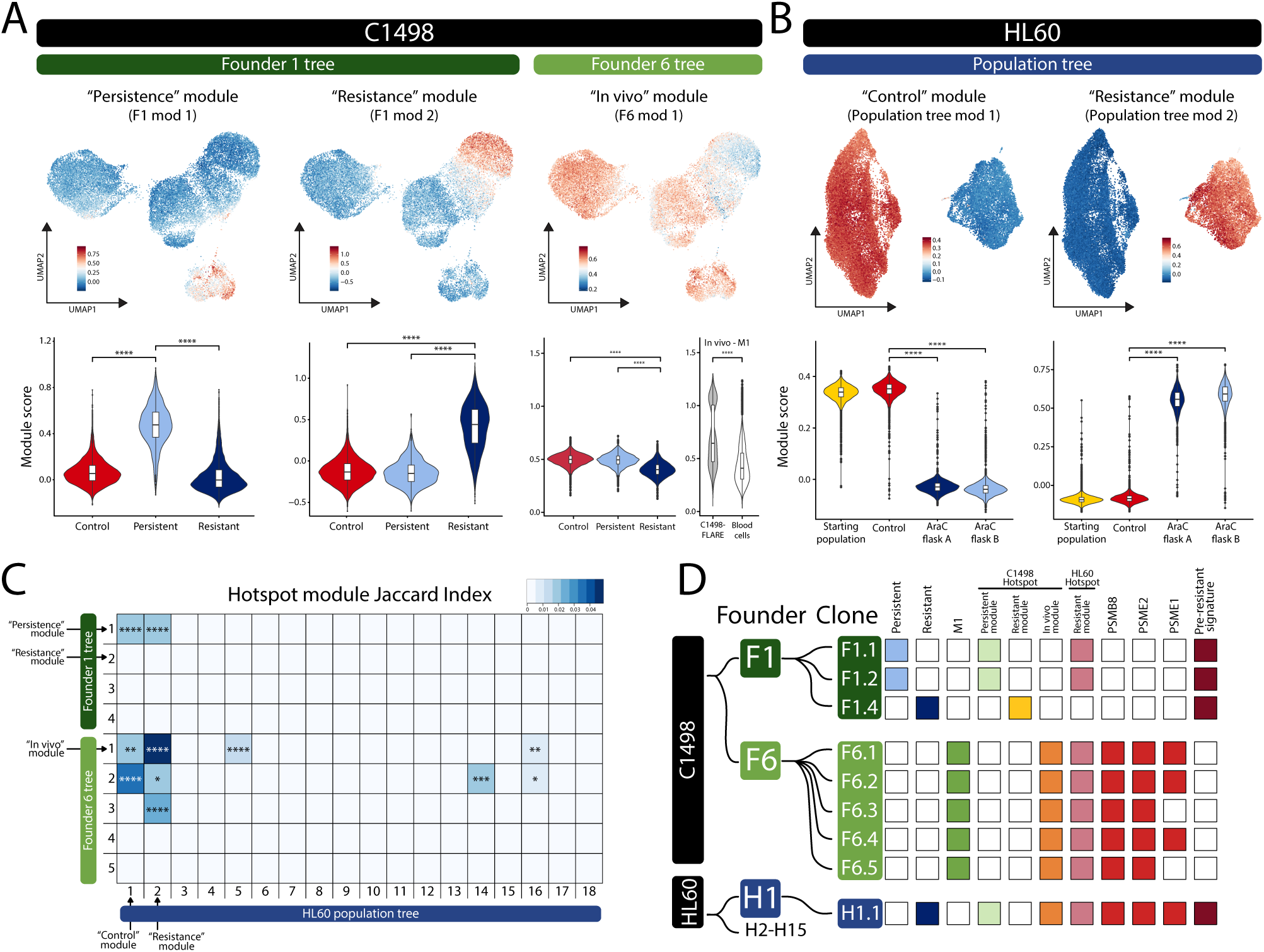
Hotspot gene modules reveal heritable AraC response mechanisms common to mouse and human AML cells. (**A**) Feature and violin plots for selected C1498-FLARE Hotspot modules. Significance determined by wilcoxon rank sum test with Bonferroni correction. (**B**) Feature and violin plots for selected HL60-FLARE Hotspot modules. Significance determined by wilcoxon rank sum test with Bonferroni correction. (**C**) Jaccard Index similarity comparing C1498-FLARE and HL60-FLARE Hotspot modules. Square color indicates Jaccard Index, asterisks represent BH-corrected P-value significance els. (**D**) Schematic showing the lineage relationships of C1498-FLARE and HL60-FLARE clones. The first three columns to the right of the clone labels indicate the setting in which each clone had a survival advantage, either in the Ara-C persistent ulation (a growth phenotype we only saw in C1498-FLARE populations), resistant population, or in AraC-treated mouse M1’s peripheral blood and bone marrow. The next four columns to the right indicate modules upregulated in each clone. The next three columns indicate whether immunoproteasome gene PSMB8 and 11S regulatory cap genes PSME1 and PSME2 were significantly upregulated in the AraC treated populations. The rightmost column indicates the upregulated expression of pre-resistant signature in AraC-treated populations.

We first focused on the overlap between HL60-FLARE “resistance” module 2 and C1498-FLARE “persistence” F1 module 1 (**Table S9**), the most heritable modules enriched in treatment conditions. The HL60-FLARE “resistance” module was distinctly enriched in all AraC-treated cells (**Figure 4B**), while the C1498-FLARE “persistence” module was associated with AraC-persistent populations (**Figure 4A**). The HL60 resistance module was primarily expressed in the H1 founder population and contained many genes found to be distinctly upregulated in treatment-naive H1 cells (**Table S6-7**). This suggests that the “resistance” module describes a lineage-specific profile that can identify cells most likely to become refractory to AraC treatment.

The HL60-FLARE “resistance” module shared 9 genes with the C1498 “persistence” module: ITSN1, VIM, CRIP1, S100A11, ANXA2, MYL6, S100A4, PSD3, and DOCK10. This set of genes is highly active in cytoskeleton maintenance and remodeling. Such pathways are implicated in the epithelial-to-mesenchymal transition (EMT), a type of transdifferentiation involved in cancer cell migration, invasion, and drug resistance^35^. While EMT is less well studied in hematological malignancies than solid tumors, there is increasing evidence of its relevance in diseases like AML^36–40^. There was no significant correlation of module expression with any tested EMT gene sets, suggesting that F1 module 1 and HL60 module 2 describe cell adhesion and motility processes largely independent from canonical EMT (**Figure S4D**). The co-upregulation and significant heritability of these genes in both cell lines under AraC treatment supports the pro-survival role for cytoskeletal dynamics separate from EMT processes, potentially constituting a form of CAM-DR. This is further supported by cell adhesion molecule genes present in the HL60 “resistance” module, including ITGB1, a primary mediator of CAM-DR^41^ and LGALS1, implicated in CAM-DR in multiple myeloma ^42^.

To identify whether heritable upregulation of CAM-DR genes in leukemic lineages prior to treatment was associated with the generation of AraC resistance, we examined related pathways in pre-experiment and control H1 cells. Sixty-five genes significantly upregulated in treatment-naive H1 cells were also found in the HL60 population tree “resistance” module, constituting an H1-specific pre-resistant state (**Figure S4E**).The majority of these genes are also involved in cell motility and adhesion, such as NCAM1^43^, PRICKLE1^44^, BMP2^45^, CPNE8^46^, and NAV3^47^. Also included are ANXA2 and PSD3, genes from the C1498 “persistence” module. While H1 cells uniquely express this pre-resistant signature prior to AraC exposure, treatment increased H1 population expression in a time-dependent manner (**Figure S4F**). In the control setting, this signature does not clearly contribute to cell fitness, but under AraC, it confers a large survival benefit.

### Identification of pro-leukemic transcriptional programs associated with in vivo immune escape

The mouse M1 “in vivo” module was the most heritable module over the F6 phylogenetic tree; it showed significant enrichment in cells collected from M1 and may represent an in vivo mechanism of survival through AraC treatment (**Figure 4A, S4G**). To determine shared mechanisms with our HL60-FLARE modules, we again compared modules and found a strong Jaccard index score between the M1 “in vivo” and our HL60-FLARE “resistance” modules (**Table S10**, **Figures 4C, S4B-C**). These two modules share immunoproteasome genes PSME1, PSME2, PSMB8, and PSMB9, immunoproteasome components involved in antigen processing and presentation for MHC class I-mediated immune response. PSMB8 and PSMB9 encode two of the three beta core subunits of the 20S immunoproteasome core complex^48^. In a variation of the immunoproteasome, the 20S core can complex with the 11S (or PA28ab) regulatory cap, the primary subunits of which are encoded by the PSME1 (PA28a) and PSME2 (PA28b) genes^49^. There is evidence that peptide fragments produced by the 20S/11S immunoproteasome may be less conducive to MHC class I presentation and lead to tumor cell immune escape^50–52^.

We investigated Psme1 and Psme2 expression patterns in F6 cells that survived engraftment and treatment in our in vivo model. Differential gene expression analysis comparing F1 and F6 in vitro revealed universal F6-specific upregulation of 11S regulatory cap genes Psme1 and Psme2 (**Table S2**). Additionally, Psmb8, Psme1, and Psme2 were significantly upregulated in F6 clones in vivo compared to their in vitro counterparts proportional to clone success in vivo (**Figures 4D, S4H**). F6 clones 1-5 upregulated Psme2 and Psmb8 in the in vivo setting, representing a founder-specific enrichment. Expression of Psme1 differentiated F6 clones further; significant in vivo upregulation was unique to clones F6.1, F6.2, and F6.4, the three most successful clones in the blood (**Figures 4D, S3B**). While Psmb8 was significantly heritable for F6 clones 1-5, Psme2 was only heritable in clones 1-3. In contrast, Psme1 expression was only heritable over the lineage tree within clone F6.5. The evolutionary plasticity of Psme1 expression in the F6.1, F6.2, and F6.4 lineages may provide a competitive advantage within the broader F6 immunoproteasome resistance mechanism.

### Patient analysis

Phylogenetic reconstruction of in vitro and in vivo AraC-treated C1498-FLARE and HL60-FLARE cells revealed heritable gene modules associated with generating and maintaining AraC resistance. To validate their clinical relevance, we used gene expression data from the TARGET AML cohort (2,007 patients, ages 0-29), the most extensive, publicly available AML patient dataset^53^. The prognostic impact of module expression in bone marrow samples, collected at initial diagnosis, showed significantly increased hazard ratios associated with the expression of HL60-FLARE “resistance” module (HR 4.99), C1498-FLARE “persistence” module (HR 2.69), and “in vivo” module (HR 3.18) (**Figure 5A**). Kaplan Meier curves confirmed that heightened expression of these modules was associated with shorter overall survival (**Figure 5B-D**). These results support the ability of FLARE lineage tracing to generate heritability patterns relevant to AML patient survival.

**Figure 5:**
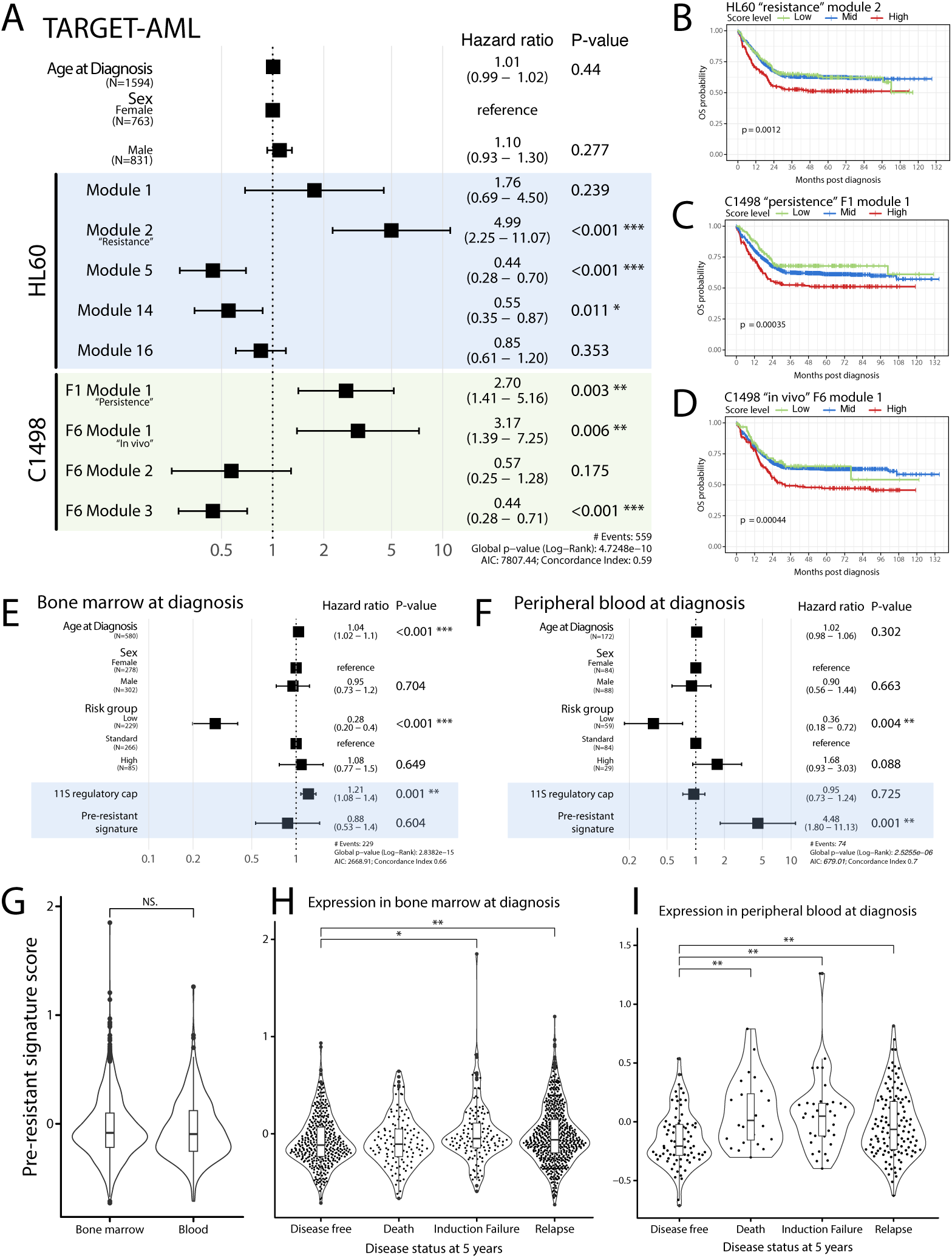
Expression levels of HL60-FLARE modules, C1498-FLARE modules, 11S regulatory cap genes, and pre-sistant signature genes are significantly associated with survival in pediatric AML patients. (**A**) Cox proportional zard ratios for HL60-FLARE and C1498-FLARE Hotspot module scores calculated from bone marrow samples taken at agnosis in the TARGET-AML cohort. (**B**) Kaplan Meier survival plots for the TARGET-AML cohort stratified into low, mid, and gh expression of HL60-FLARE “resistance” module 2. Corresponding risk table can be found in supplementary table S11. (**C**) plan Meier survival plots for the TARGET-AML cohort stratified into low, mid, and high expression of C1498-FLARE ersistence” F1 module 1. Corresponding risk table can be found in supplementary table S11. (**D**) Kaplan Meier survival plots the TARGET-AML cohort stratified into low, mid, and high expression of C1498-FLARE “in vivo” F6 module 1. rresponding risk table can be found in supplementary table S11. (**E**) Cox proportional hazard ratios for 11S regulatory cap d pre-resistant scores calculated from bone marrow samples taken at diagnosis in the TARGET-AML cohort. (**F**) Cox oportional hazard ratios for 11S regulatory cap and pre-resistant scores calculated from peripheral blood samples taken at agnosis in the TARGET-AML cohort. (**G**) Violin plot showing pre-resistant signature scores calculated from samples taken at agnosis. Bone marrow and blood samples do not significantly differ in signature scores. (**H**) Violin plot showing pre-resistant nature scores calculated from bone marrow samples taken at diagnosis. Mean scores for patients who experience induction lure or relapse within 5 years post diagnosis are minimally but significantly higher than patients who are disease free 5 years post diagnosis. Significance was calculated with Wilcoxon rank sum tests and Bonferroni corrected. (**I**) Violin plot showing pre-sistant signature scores calculated from peripheral blood samples taken at diagnosis. Mean scores for patients who experience death, induction failure, or relapse within 5 years post diagnosis are significantly higher than patients who are disease free 5 years post diagnosis. Significance was calculated with Wilcoxon rank sum tests and Bonferroni corrected.

We then focused on our pre-resistant/CAM-DR sub-module and 11S cap subunits of the 20S/11S immunoproteasome variant to explore their roles in treatment response and survival. We generated Cox proportional hazard ratios, including patient risk groups as covariates to account for interactions between risk group and signature expression. Expression levels of both signatures were associated with the risk of death independent of predefined risk groups. Interestingly, signature-associated risk of death differed drastically when patient expression scores were calculated from bone marrow or peripheral blood taken at diagnosis. In bone marrow samples, expression of 11S regulatory cap genes, but not the pre-resistant signature, was significantly associated with the risk of death (HR 1.21, p-value < 0.001, **Figure 5E**). Conversely, in peripheral blood, expression of the pre-resistant signature, but not the 11S regulatory cap genes, was significantly associated with a greater risk of death (HR 4.48, p-value < 0.001, **Figure 5F**). While 11S regulatory cap genes were expressed slightly more in bone marrow samples, potentially explaining why PSME1 and PSME2 expression have higher prognostic impact in bone marrow, there was no significant difference in expression of the pre-resistant sub-module between sample types (**Figure S5A, 5G**).

We then assessed the relationship between signature expression in pre-treatment diagnosis samples and patient status at 5 years post-inclusion. Specifically, we compared expression levels between patients who experienced death, induction failure, or relapse within 5 years compared to patients who remained disease-free for 5 years. Patients who experienced induction failure or relapse, but not death, tended to have higher pre-resistant sub-module scores in their blood and bone marrow samples at diagnosis (**Figure 5H-I**). Importantly, these trends were more dramatic when comparing blood samples at diagnosis; patients experiencing induction failure, relapse, and death expressed significantly higher levels of the pre-resistant signature in blood compared to patients disease-free at 5 years. There was a slight negative relationship between pre-resistant sub-module score and peripheral blast percentage at diagnosis, suggesting this pattern does not reflect peripheral leukemic burden (**Figure S5B-C**).

## Discussion

Chemotherapy resistance likely arises from the selection of pre-existing resistant populations and the reprogramming of previously sensitive cells. There is an unmet need for tools to dissect this evolution over time. We developed FLARE, a CRISPR-based dynamic lineage tracing system capable of tracking founder, clone, and subclone lineage in tandem with single-cell transcriptomics in response to chemotherapy. We demonstrated how our hierarchical approach can record cancer transcriptional evolution over an extended treatment time course, capturing both clonal evolution and selection of pre-existing cellular states. We used this technology to dissect the transcriptional evolution of AML, leveraging both in vitro and vivo systems and mouse and human models.

Using FLARE we uncovered the (1) heritable patterns of cell adhesion-associated AraC-tolerance and the (2) upregulation of immunoproteasome 11S regulatory cap subunits contributing to clone-specific in vivo fitness. The expression of both signatures in AML patients is significantly associated with shorter overall survival. High expression of PSME1 and PSME2, the major subunits of the 11S regulatory cap, in bone marrow but not peripheral blood, is associated with poorer prognosis. In contrast, high expression of our pre-resistant signature, which consists of various genes involved in cytoskeletal and cell adhesion pathways, is more informative when measured in blood samples taken at diagnosis. Both the 11S regulatory cap and pre-resistant signatures are composed of many genes not previously implicated in AML.

Our pre-resistant signature comprises many cell adhesion-mediated drug resistance (CAM-DR) genes. This mechanism has long been studied in the context of solid tumors, but its importance in hematological malignancies is growing^54,55^. Leukemic CAM-DR is thought to result from interactions between AML blasts and bone marrow stromal cells via various integrin molecules^56^. These interactions may protect leukemia cells, leading to minimum residual disease persistence and treatment failure upon disease recurrence^55^. This increased survival is thought to result from anoikis resistance, in which cells avoid apoptosis in response to becoming detached from the extracellular matrix^57^. Interestingly, the emergence of CAM-DR in our work was primarily seen in cells treated with AraC in standard in vitro cell culture. This may indicate that treatment alone can lead to such resistance outside of the bone marrow niche.

Furthermore, survival analysis of the TARGET AML cohort revealed that expression of our potential CAM-DR-associated pre-resistant signature had a significantly higher prognostic impact when measured from peripheral blood samples taken at diagnosis. Our survival analysis also suggests that the presence of leukemic blasts with enrichment of cytoskeletal, cell adhesion, and cell motility processes in the blood before induction therapy is associated with induction failure, death and relapse within 5 years of diagnosis. Therefore, patients with “pre-resistant” lineages prior to treatment are less likely to benefit from induction chemotherapy, with cells already displaying CAM-DR that can survive treatment and expand to outcompete other leukemic lineages. This mechanism needs further exploration, but it suggests a secondary role of CAM-DR in AML outside of the bone marrow niche.

While the exact pro-leukemic mechanism of our 11S signature cap is still unclear, there are data to suggest the 11S regulatory cap leads to the generation of fewer peptides conducive to MHC class I presentation and can interfere with tumor-specific epitope production^51,52,58^. 11S subunit expression levels in pediatric AML patient bone marrow samples taken at diagnosis are significantly associated with an increased risk of death and shorter overall survival. PSME1 and PSME2 expression levels in bone marrow samples differed negligibly between patients with adverse outcomes within 5 years compared to those that remained disease-free. This indicates that while PSME1 and PSME2 expression in bone marrow is associated with a higher risk of death, it is not highly predictive of induction failure or relapse within 5 years of diagnosis. It may instead indicate the presence of leukemic lineages that are better able to evade immune detection, resulting in a more aggressive and “colder” immune microenvironment.

In the future, the FLARE approach could be used to dissect the transcriptional evolution of other tumor types under selective treatment pressure. Given the ability to sample tumor evolution from tagged cell populations in circulating blood, this can be multiplexed to explore many more initial cell states and treatment spaces. Moreover, this approach can be coupled to mouse lineage-tracing models such as DARLIN^59^ or the MARC1^60^ mouse to capture the interplay between engrafted tumor and immune cell populations in the context of checkpoint inhibitors and other immunotherapies.

## Methods

### Cell Lines and drugs

The murine C1498 cell line (Cat# TIB-49) was purchased from ATCC in 2021. The HL60 AML cell line was a gift from Dr. Bonnie Lau’s lab. C1498 cells were cultured in DMEM, supplemented with 10% FBS, 100 u/mL penicillin, and 100 ug/mL streptomycin. HL60 cells were cultured in Iscove’s Modified Dulbecco’s Medium, supplemented with 20% FBS, 100u/mL penicillin, and 100ug/mL streptomycin. All cell lines were expanded, aliquoted, and cryopreserved in culture media plus 5% DMSO and stored in liquid nitrogen upon receival.

Active cell cultures were stored at 37°C with 5% CO_2_. All cultured lines were regularly checked for the presence of mycoplasma using the Universal Mycoplasma Detection Kit (ATCC, 30-1012K). Cytarabine (AraC) powder was purchased from Selleck Chemicals (Cat # S1648). AraC was resuspended in PBS fresh for each in vitro and in vivo treatment.

### Mice

KH2/iCas9 mice (Jackson Laboratory strain #029415) were a gift of the Huang lab at Dartmouth College. All mice were maintained in pathogen-free housing at the Center for Comparative Medicine and Research in Dartmouth Hitchcock Medical Center. All experimental protocols were approved by and performed in accordance with the guidelines of the Institutional Animal Care and Use Committee at Dartmouth College. To attenuate the immune response to Cas9, all injected KH2/iCas9 were exposed to doxycycline during in utero development through irradiated feed containing 625ppm doxycycline (ScottPharma, Inc. Marlborough, MA).

### Molecular cloning

#### GESTALT lineage tracing system engineering

All digested vectors and fragments and PCR products were purified with the Monarch Gel Purification Kit (Cat# T1120S) or the Zymo Clean and Concentrate-5 Kit (Cat# D4003). All Gibson assembly reactions were conducted with NEB HiFi DNA Assembly 2x Master Mix (Cat# E2621S), following the ssDNA HiFi assembly protocol. Golden Gate assemblies were conducted following the Rice University Esp3I protocol. Chemical bacterial transformations were done with DH5alpha competent cells following the NEB Transformation Protocol with a 2-hour recovery at 37°C. Electroporated transformations were conducted with NEB Electrocompetent E. coli and were electroporated with the following conditions: 2kV, 200Ohms, 25uFd. All bacterial cultures were grown in a 37°C shaker at 220-290rpm overnight. Vectors were purified using either the Qiagen Miniprep Kit or the Zymo Midiprep II Kit. Correct cloning of all final constructs was confirmed via Circuit-seq^61^.

#### FLARE-miniV10-10x-UMI

Eight Cas9 target sequences were designed for high efficiency (>69% Doench) and low off target scores. The eight-target array was purchased as a gene block from IDT, with upstream Esp3I sites for founder tag insertion and NdeI + BamHI sites for cloning purposes. The target gBlock and pLARRY-EGFP vector (Addgene #140025) were digested with NdeI+BamHI and ligated together with Quick Ligase (Cat# M2200s). Resulting construct is referred to as pLARRY-targets.

The PB-CMV-MCS-EF1-puro vector (Systems Bio cat #PB510B-1) was digested with SpeI and HincII, removing the puromycin resistance gene, and then circularized with a Gibson assembly reaction with a short single stranded joining oligo. The resulting construct is referred to as miniPB.

The FLARE-miniV10-10x vector was then constructed by amplifying EGFP and the target array from pLARRY-targets. The PCR primers used KpnI and EcoRI sites. The amplified EGFP+target array and miniPB were digested with KpnI and EcoRI, gel purified, and ligated together using T7 ligase. The 10X capture sequence 1 was incorporated via Gibson assembly. The FLARE-miniV10-10x construct is available at Addgene (Plasmid #234492).

Founder tag oligos consisting of 14 degenerate bases (the founder tag) and Esp3I sites were purchased from IDT. One cycle of PCR was used to anneal and extend the oligos. Founder tag oligos were then inserted into FLARE-miniV10 via a Golden Gate Assembly reaction; the resulting product is FLARE-miniV10-10X-UMI.

#### FLARE-pbV10g-neo

Individual Cas9 guides complementary to the target sequences in FLARE-miniV10 were designed and cloned into the pX330-mCherry (Addgene #98750) vector using the Zheng Lab pX330 cloning protocol. Guides, along with their individual U6 promoters, were amplified from pX330-mCherry vectors. A secondary PCR reaction added flanking Esp3I sites to each U6-guide fragment. Esp3I sites were also added to the PB-CMV-MCS-EF1-puro vector, while the puro resistance gene was removed. U6/guide sequences were joined and inserted into PB-CMV-MCS-EF1-puro via a 9-part Golden Gate Assembly reaction. The neomycin resistance gene was amplified from the hCas9 vector and inserted into the U6/guide array vector via Golden Gate Assembly. The FLARE-pbV10g-neo construct is available at Addgene (Plasmid #234493).

In earlier versions of FLARE constructs, Cas9 guides were encoded into a single construct along with the lineage recorder Cas9 target array. For reasons not yet fully understood, splitting this into two vectors (FLARE-miniV10-10X and FLARE-pbV10g-neo) led to significantly higher editing rates across targets.

#### FLARE-pbCas9-tag-mCherry

The miniPB construct was digested with KpnI for a Gibson Assembly insertion of a landing pad containing a NotI and a new KpnI site. The resulting vector is referred to as miniPB-LP. The pX330 construct was digested with KpnI and NotI to isolate the Cas9 transgene, which was then ligated into the landing pad miniPB-LP vector. The resulting construct is referred to as miniPB-Cas9.

The pX330-mCherry construct was digested with BamHI and HindIII for the insertion of a clone tag landing pad containing type IIs restriction sites in the 3’ UTR of the mCherry transgene using Gibson Assembly. The mCherry transgene, including the clone tag landing pad, was amplified and extended to include Gibson Assembly adaptors complementary to the insertion region in miniPB-Cas9. The miniPB-Cas9 construct was then digested with NotI and the mCherry and clone tag landing pad were inserted using Quick Ligase. The resulting construct is referred to as FLARE-pbCas9-mCherry, and is available at Addgene (Plasmid #234494).

Lastly, clone tag oligos with Esp3I sites were purchased from IDT. For C1498-FLARE cells, the clone tag consisted of 14 degenerate bases followed by 10x capture sequence 2. For HL60-FLARE cells, the clone tag consisted of 38 patterned bases (NNNWSNNNWSNNNWSNNNWSNNNWSNNNWSNNNWSNNN, pattern from Johnson et al ^62^) followed by 10x capture sequence 1. One cycle of PCR was used to anneal and extend the oligos, and the resulting product was inserted into FLARE-pbCas9-mCherry via Golden Gate Assembly. Capture sequence 2 was replaced by capture sequence 1 for HL60-FLARE cells due to the planned discontinuation of support for capture sequence 2 announced by 10x.

### Cell engineering and lineage tracing system testing

We used the Neon Electroporation System to integrate all piggyBac constructs into cells to generate stable cell lines. Electroporations were carried out with the following protocols: C1498 cells were resuspended at 2×10^7 cells/mL and electroporated with 1080V, 50ms, and 1 pulse; HL-60 cells were resuspended at 2×10^7 cells/mL and electroporated with 1450V, 35ms, and 1 pulse. All electroporations were done with the Neon 100ul tip system. Transposase:transposon ratios for piggyBac integration ranged from 1:1 to 1:8. All FACS was conducted on a BD FACSAria II or BD FACSAria Fusion. For antibiotic selection cells were incubated with selection media for 1-3 weeks, and dead cells were removed from culture 2-3 times/week using a Ficoll density gradient.

GESTALT constructs were introduced to cell lines in the following order: first low passage cells were electroporated with FLARE-miniV10-10X-UMI and FLARE-pbV10g-neo. EGFP-positive and neomycin resistant cells were isolated via FACS and 1ug/mL G418 selection. This step was repeated two to three more times, gating for higher EGFP expression during FACS in each subsequent round. For the third round of FACS for the C1498 cell line, individual cells were sorted into wells of a 96-well plate to initiate individual clonal populations.

Prior to lineage tracing experiments, engineered lineage tracing cells were electroporated with FLARE-pbCas9-tag-mCherry and mCherry+ cells were isolated via FACS. Cells were sorted into bulk populations and 96-well plates at 100 cells/well.

Lineage recorder editing levels in C1498-FLARE cells over time were calculated using single cell and bulk lineage barcode data. Day 0 editing was calculated from bulk (not a subset of founders) C1498-FLARE cells prior to FLARE-pbCas9-tag-mCherry integration; day 13 editing was calculated from this same population 13 days following integration post FACS. The remaining editing rate timepoints were measured using the cell population used in C1498-FLARE founder pool AraC treatment experiment and lineage recorder reads were subset for reads containing valid founder tags associated with founders 1, 5, 6, and 11. Day 44 editing was calculated from single cell sequencing of the founder pool subset for F1 and F6 prior to AraC treatment. Day 64 editing was calculated from single cell sequencing of one untreated flask cultured from the same population as used in day 44, and was subset for valid cells identified via GEX analysis. Day 87 was also measured from one control flask from this population – editing was calculated from single cell lineage endpoint data.

### C1498-FLARE founder isolation and characterization

To screen individual C1498-FLARE founder populations for copies of FLARE-miniV10-10X-UMI integrations gDNA was collected and founder tags were amplified and prepared into sequencing libraries. Founder tag libraries for individual populations were multiplexed and sequenced on an Illumina MiniSeq with a Mid-Ouput 300-cycle kit. Founder tags for each clone were determined as having > 4,300 reads. Cells from each clone were expanded, aliquoted, and cryopreserved.

#### Growth curves

To begin growth curves for individual founders, cells were resuspended at 50k cells/mL and plated in T25 flasks in triplicate. From day 0-7, two 10ul aliquots from each flask were mixed with 10ul Trypan Blue and counted on the Luna II Cell Counter daily. Growth curve data were analyzed and plotting using the Growthcurver R package ^63^.

#### AraC dose-response curves

Cells were plated at 15,000 cells/well in a 96-well plate in media containing various AraC dosages in triplicate (0.0001mg/ml-10mg/ml). 100ul of CellTiter 96 AQueous One Solution (Cat# G3582) was added to each well and after incubation for 1-3 hours absorbance was measured at 490nm on a SpectraMax i3x. Absorbance from wells containing media without cells or drug were averaged to calculate background absorbance. Corrected absorbance for each well was calculated as (well absorbance – background absorbance). Corrected absorbances were then loaded into the drc R package for generating dose-response curves, ED50 values, and EC50 values^64^.

#### Whole genome sequencing

High molecular weight (HMW) gDNA was isolated from parental C1498 cells and individually engineered clones using the Monarch High Molecular Weight gDNA Kit for Cells (Cat# T3050S). HMW gDNA samples were multiplexed and prepared into a sequencing library following the Oxford Nanopore SQK-NBD114.24 kit. The library was then sequenced on a PromethION flow cell with R10.4.1 chemistry. Sequencing data were basecalled and demultiplexed using Guppy. Variants were called using PEPPER-Margin-DeepVariant, SNPs were isolated and filtered for a minimum quality of 20 and mean depth between 10 and 80. To remove population variants, the GRCm39 dbSNP was annotated using Ensembl VEP and variants with moderate or high impact were removed. The resulting dbSNP was subtracted from filtered SNPs using Bedtools ^65^.

Remaining variants were annotated with Ensembl VEP and intersected with datasets of known AML mutations.

## AraC response and resistance experiments

### C1498-FLARE founder pool treatment response and resistance

C1498-FLARE founders F1, F5, F6, and F11 were sorted by FACS on day 13 following electroporation with the FLARE-pbCas9-tag-mCherry vector. Cells were sorted into 96-well plates at 100 cells/well. In vitro AraC treatment began on day 44 (31 days post FACS). Cells were allowed to grow for a longer period following sorting due to growth rate differences between founders. On day 44, 1 well of sorted cells for each clone was pooled at equal numbers and split into four flasks for in vitro treatment with an aliquot set aside for mouse injections.

#### In vitro

Two flasks were maintained in normal culture media, the remaining two flasks were maintained in media containing 0.05ug/ml AraC. Flasks were maintained for 43 days to allow for the development of AraC resistance. Media was replaced 2-3 times/week and split when needed. Dose-response curves were calculated for each flask on day 15 of treatment, and a final dose-response curve was calculated for flask displaying resistance on day 36 of treatment. On day 43 of treatment, cells were prepared for MULTIseq tagging and single cell capture following MULTIseq and 10x protocols.

#### In vivo

Pooled C1498 clone cells were washed twice with PBS, resuspended at 250,000 cells/mL and filtered through a 40uM filter. Mice were injected IV with 100ul of cell suspension (25,000 cells/mouse). On day 14 mice were randomly assigned to treatment groups; on days 14-16 treatment mice received 100mg/kg AraC IP and control mice received an equal volume of PBS. Blood was collected via the retro-orbital sinus on days 13, 28, 43, 50, and 57. Prior to blood draws mice were anesthetized with Isoflurane. Microhematocrit capillary tubes were placed in 1.7mL tubes containing 50mM EDTA and then used to collect 50-200ul of blood from the retro-orbital sinus into EDTA coated tubes. Blood samples were then kept on ice until further processing. Mice displaying symptoms of high AML burden were euthanized. All surviving mice were euthanized on day 67 post cell injections. Upon euthanasia whole blood was collected via cardiac puncture and femoral bone marrow was harvested. gDNA was isolated from 20ul of whole blood, and remaining blood was separated using a Ficoll-paque density gradient to isolate PBMCs. To isolate bone marrow, 18g needles were used to puncture the bottom of 0.5mL tubes, femur bones were placed in the 0.5mL tubes and nested in 1.7mL tubes, and tubes were spun at > 10,000g for 20 seconds. Bone marrow pellets and PBMCs were methanol fixed and permeabilized following 10x protocols.

#### Human AML cell line in vitro treatment response

Engineered lineage tracing HL60-FLARE cells were sorted into 96-well plates at 100 cells/well following electroporation with FLARE-pbCas9-tag-mCherry to establish GESTALT lineage tracing populations, as described above. Sorted well populations were expanded for 23 days. On day 23 post FACS, five expanded wells were combined into an initial cell pool. The pool was then split for 10x scRNA-seq preparation, methanol fixation, cryopreservation, and in vitro AraC treatment. For in vitro AraC treatment cell pools were split into control (no AraC) and AraC conditions (0.01ug/mL AraC). All treatment wells were cultured in duplicates. AraC solutions and media were made fresh for media changes twice per week, and wells were split as needed. Cell concentration for each well was measured once per week in duplicates using the Luna II Cell Counter. Cells were regularly cryopreserved through the experiment for later analysis. In vitro treatment continued for 89 days, after which cells were prepared for 10x scRNA-seq preparation with MULTIseq multiplexing.

## Lineage library preparation and sequencing

All PCR reactions were conducted using Q5 HotStart Polymerase (Cat # M0494L) and Sybr Green dye. DNA was purified from PCR reactions using Omega TotalPure NGS magnetic beads with 0.8-1.5x ratios based on amplicon size.

### Bulk DNA library preparation and sequencing

gDNA was collected from cultured cells, PBMCs, and bone marrow samples using the Qiagen DNeasy Blood and Tissue Kit (Cat# 69504). For library preparation, lineage recorders (including founder tags and clone tags) were first tagged with sequencing UMIs prior to amplification to control for PCR bias, with the exception of libraries constructed from blood collections. Blood collection libraries were treated as sensitive samples, as C1498-FLARE cells were sparse in the blood. As a result, these libraries were amplified without sequencing UMIs and initial PCRs were set up in a PCR hood to prevent contamination. For all libraries, TruSeq/Nextera indices and Illumina adaptors were subsequently attached with PCR. Final sequencing libraries were quantified with the Qubit fluorometer and the NEB NextQuant Library Quant kit (Cat # E7630L). Libraries were then diluted to the appropriate loading concentration for sequencing.

### Single cell library preparation and sequencing

Cells were captured for single cell sequencing following 10x guidelines for 3’ scRNA-seq with feature barcoding. For multiplexed single cell experiments, cells were first tagged following MULTIseq protocols (Millipore Sigma, LMO001)^66^. Gene expression and MULTIseq libraries were constructed and quantified following 10x and MULTIseq guidelines with one alteration – a portion of cDNA was set aside following initial cDNA amplification but prior to SPRIselect cleanup to be used for FLARE lineage library construction. This was done to avoid splitting FLARE transcripts between GEX and feature barcoding fractions.

The previously set aside cDNA was purified using a 2x bead cleanup. For each sample, cDNA was split to generate a library of dynamic lineage recorders (which includes founder tags) and a library of clone tags. All single cell FLARE libraries were initially amplified in 9-16 replicate wells and then combined into 3-4 separate sub-libraries. TruSeq/Nextera indices and Illumina adaptors were subsequently attached with PCR. Final sequencing libraries were quantified with the Qubit fluorometer and the NEB NextQuant Library Quant kit (Cat # E7630L). Libraries were then diluted to the appropriate loading concentration for sequencing.

## Computational analysis

### Sequencing data pre-processing

For bulk DNA sequencing raw data were converted to fastq format using bcl2fastq2 or BCL Convert (Illumina). For single cell RNA sequencing raw data were converted to fastq format using CellRanger mkfastq. Cells were then called using CellRanger count for non-multiplexed 10x runs or CellRanger multi for MULTIseq multiplexed 10x runs^67^.

### Single cell RNA-seq analysis

Cellranger output filtered matrices were analyzed with Seurat 5.1.0^68^. Low quality cells were filtered out based on the number of RNA features and percentage of mitochondrial reads. For non-multiplexed samples doublets were identified and removed using the scDblFinder R package^69^. For multiplexed samples, MULTIseq barcode filtered matrices were added as Assay Objects to Seurat objects. Cells were demultiplexed and doublets were identified and removed with the *MULTIseqDemux()* function. For PBMC and bone marrow samples cell type annotations were added using the SingleR R package with the MouseRNAseqData reference set from the celldex R package^70^. All single cell Seurat objects were normalized with the SCTransform R package for data visualization and integration with other objects^71,72^. Differential gene expression analysis was conducted using the *FindMarkers()* and *FindAllMarkers()* Seurat functions.

### Gene set enrichment analysis

C1498-FLARE founder-specific gene set enrichment analysis was conducted with the fGSEA R package^73^. Differentially expressed genes were identified for each founder by Wilcoxon rank sum test and input into the fGSEA function with Mus musculus Hallmark gene sets from mSigDB.

### Single cell lineage analysis and tree reconstruction

#### Founder and clone tag clustering

To map founder tags to founder tag groups (or founders) and clone tags to clone tag groups (or clones), we used a clustering strategy based on per cell tag set similarity. Briefly, given a mapping of cells to a set of tags, a graph was constructed with cells as nodes and Jaccard similarity as edges. Jaccard similarity was calculated pairwise as the intersection between per cell tag sets divided by the union of per cell tag sets. Any pairs of cells with a Jaccard similarity of 0 (indicating no overlapping tags) shared no edges. Cells were assigned to clusters using the Louvain algorithm for community detection which returned the partition that maximized modularity and with resolution = 1.

#### Founder assignment

To extract lineage information from scRNA-seq-captured dynamic lineage recorders, we first used SingleCellLineage, as previously described^13,14^. Briefly, sequences are collapsed by 10x UMI and aligned to the reference to obtain a pattern of insertions and deletions across a target array per cell. Founder tags, located immediately upstream of the lineage recorders, were extracted per collapsed merged read. We filtered cell barcodes against the 10x feature barcoding Cell ID whitelist. Merged reads with less than 85% identity to the reference sequence were removed from the dataset. To ensure we were using high quality barcodes to build single-cell lineage trees, we leveraged tools from the Cassiopeia pipeline^74^ to perform additional data cleaning. Founder tags were corrected for PCR errors via Starcode^75^ with Levenshtein distance = 1. UMIs were error corrected using Cassiopeia function *error_correct_umi()*. Finally, we filtered our error corrected founder tags and target arrays using *filter_molecule_table()* to remove low read count UMIs and cells with too few UMIs. Additionally, this step ensures there is only one target array per 10x cell ID-founder tag combination.

Founders were assigned by the combination of founder tag integrations in cells. For C1498-FLARE founder populations, founder tag assignments were previously determined via sub-founder culturing and bulk gDNA sequencing. Therefore, if cells had reads mapping to more than one founder based on the founder tags found in its lineage data, it was assigned to a founder based on the total sum of reads and UMIs associated with a certain founder group. Only cells with 90% identity to a certain founder were kept.

Unlike the C1498-FLARE founders, HL60-FLARE founders had no ground truth mapping of founder tags to founder groups. Subsequently, we took additional steps to ensure we had a confident list of founder tags per cell and group. To remove within-cell background noise, we removed integrations that made up less than 1.5% of reads per cell. This threshold was based on qualitative inspection of the distribution of integration read counts per integration per cell.

The founder tags were further filtered by removing tags found in fewer than 5 cells across the dataset. The cleaned set of tags were clustered into founders as described previously (*Founder and clone tag clustering*). Finally, to refine the final set of founder tags defining a founder, we filtered for tags with greater than 50% identity to a single founder group. This threshold was based on visual inspection of the distribution of tag identity to founders, determined by the total number of cells with a tag in a founder divided by the total number of cells with a tag. This additionally ensures that each founder tag is found in one founder.

#### Clone assignment

Clone tags were extracted from sequencing reads using the SingleCellLineage pipeline. For HL60-FLARE clone tag reads, only tags matching the expected 38bp IUPAC pattern were kept. Clone tag reads with more than one missing base within the tag sequence window were removed from the dataset, then clone tags were error corrected using Starcode with Levenshtein distance = 1 for 14 bp tags or Levenshtein distance = 2 for 38 bp tags. These clone tags were then filtered for read count > 1 and number of UMIs > 1. Tags with nucleotide repeats of 5 or longer were also removed (i.e. AAAAA, TTTTT, GGGGG, CCCCC). Background tags, defined as contributing to fewer than 10% of all reads in the cell, were filtered out. Finally, clone tags were discretized to founders by keeping tags with 90% identity (C1498s, 2 founders) or 50% identity (HL60s, 15 founders) to a given founder based on the total proportion of reads per clone tags. C1498-FLARE and HL60-FLARE cells were assigned to clones as previously described (*Founder and clone tag clustering*) given the final set of clone tags.

#### Building Trees

Trees were reconstructed per clone using Cassiopeia’s^74^ greedy vanilla solver from the pattern of indels per target array. The method was chosen for its scalability given the size of our trees ranges from tens to thousands of cells. Briefly, a tree is built in a top-down fashion by splitting cells into two groups based on the most frequent mutation in the population. Cell subsets are repeatedly split until only one cell remains per subset. Clones belonging to the same founder population were combined into founder trees using a custom python script to attach trees at the root, due to the hierarchical nature of the lineage tracing tag system. All founder trees were combined with the same method to generate a population tree. Cells with unedited/wildtype target arrays were removed from the analysis prior to building the trees.

#### PATH analysis

To identify genes that were significantly heritable in our clonal population we used the recently published tool, PATH ^28^. PATH takes as input a phylogenetic tree along with either discrete annotations like cluster assignments or continuous data like gene expression or module scores. We considered the top 3000 highly variable features per tree to identify genes with strong autocorrelations, suggesting high heritability.

#### Hotspot module generation

To identify informative gene expression modules based on our single-cell lineage trees and gene expression data, we leveraged the tool Hotspot^34^ as described in the following vignette https://hotspot.readthedocs.io/en/latest/Lineage_Tutorial.html. Briefly, we ran Hotspot on our log-normalized data via the ‘normal’ model filtered for genes in a minimum of 10 cells. Then we generated the KNN graph with 10 neighbors. Autocorrelation scores are calculated per gene to perform feature selection. These genes are then correlated to generate a gene-by-gene matrix, which is used to identify modules using a hierarchical clustering approach with min_gene_threshold = 10 to control the size of modules being joined.

#### Module overlap analysis

C1498-FLARE module mouse genes were converted to human orthologs for analysis. C1498-FLARE and HL60-FLARE modules were assessed for significant overlap as measured by Jaccard indices and odds ratios compared to genomic background using the GeneOverlap R package^76^.

##### Survival analysis

TARGET-AML RNAseq, clinical, and sample data were accessed through the cBioPortal. Expression datasets were split by sample type where applicable. Sample expression scores for modules and gene sets were calculated by averaging the FPKM or RPKM scores. Cox proportional hazard ratios were generated using the survival R package^77^. To stratify patients for Kaplan-Meier analysis, we first determined the expression score mean and standard deviation within the samples of interest. Patients with scores within one standard deviation of the mean were assigned to the Mid-expression group. Patients with scores above and below one standard deviation of the mean were assigned to the High and Low expression groups, respectively. We excluded patient samples missing information for tissue type.

## Acknowledgments

We thank the members of the McKenna lab for experimental help and scientific input. We especially thank Aidan Cook for his computational expertise. In addition, we thank Prerna Malaney and Bonnie Lau for their expertise in AML, and Brock Christensen for his input on survival analyses. We acknowledge the Dartmouth Cancer Center and the shared resources supported by NCI Cancer Center Support Grant 5P30CA023108. We especially thank Fred W. Kolling IV, Laurent Perreard, Carol Ringelberg, and Elizabeth Sergison of the GMBSR for their sequencing guidance and expertise. FACS was carried out by Gary Ward in DartLab, the Immune Monitoring and Flow Cytometry Shared Resource at the Dartmouth Cancer Center (RRID: SCR_019165). We also acknowledge the Dartmouth Cancer Center Irradiation, Imaging, Microscopy, and Animal Cancer Models Shared Resource (RRID:SCR_023478), and specifically Jennifer Fields for her help with animal experiment strategy. All Illumina sequencing was carried out in the Genomics and Molecular Biology Shared Resource (RRID:SCR_021293) which is additionally supported by NIH S10 (1S10OD030242) awards. Single-cell studies were conducted through the Dartmouth Center for Quantitative Biology in collaboration with the GMBSR with support from NIGMS COBRE (P20GM130454) and NIH S10 (S10OD025235) awards. This work was supported by DP2GM149750 and the V Foundation, and A.M. is supported by The Pew Biomedical Scholars program.

## Data availability

All sequencing data associated with this publication have been deposited in NCBI’s Gene Expression Omnibus^78^ and are accessible through GEO Series accession numbers GSE289454 and GSE289459.

---

**Supplemental figure 1:**
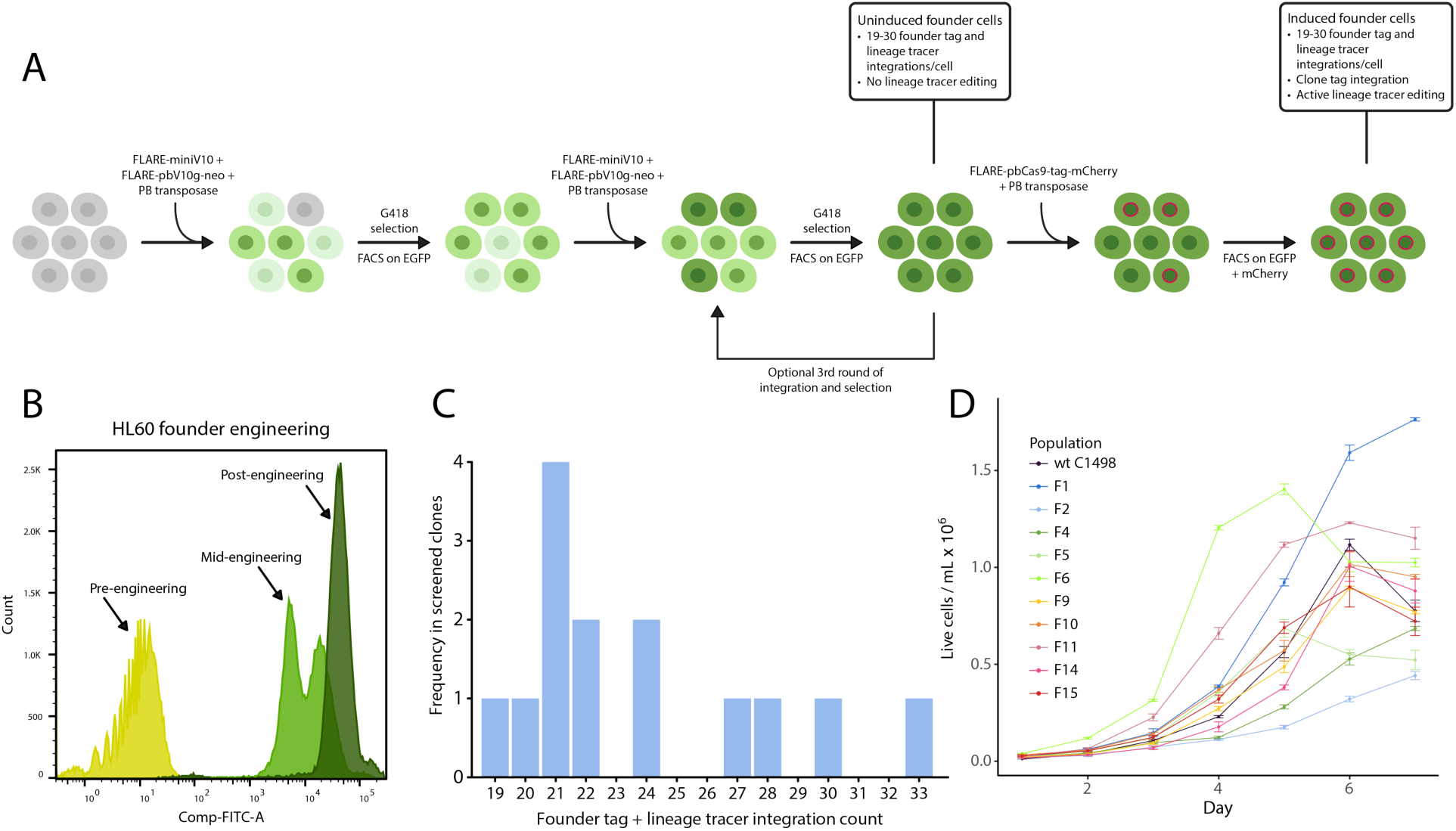
FLARE cell line engineering allows for the generation of founder populations maintaining intra-founder heterogeneity. (**A**) Schematic of engineering lineage tracing AML cells. Wildtype cells are first electroporated with FLARE-miniV10-10X-UMI + FLARE-pbV10g-neo + PB-transposase. Successfully integrated cells are selected via G418 treatment (FLARE-pbV10g-neo) and FACS for EGFP (FLARE-miniV10-10X-UMI). This process is repeated 1-2 more times, successively selecting for cells with high numbers of integrations. Once the desired number of integrations is reached, cells are considered uninduced founder cells. Prior to the start of a lineage tracing experiment, uninduced founder cells are electroporated with FLARE-pbCas9-tag-mCherry and PB-transposase and sorted on EGFP (FLARE-miniV10-10X-UMI) and mCherry (FLARE-pbCas9-tag-mCherry). Resulting cells are referred to as induced founder cells and contain all the necessary integrations for founder and clonal tracking. (**B**) FACS plots showing the increase in EGFP expression in HL60 cells through 2 rounds of electroporation with FLARE-miniV10-10X-UMI + FLARE-pbV10g-neo + PB transposase. Post-engineering uninduced HL60-FLARE founder cells uniformly express high levels of EGFP. (**C**) Uninduced C1498-FLARE founder populations were subcultured to determine founder tag and lineage recorder integration counts. Founders contained between 19 and 33 integrations. (**D**) Subcultured founder populations display a range of growth rates and carrying capacities, indicating sustained heterogeneity following the engineering process.

**Supplemental figure 2:**
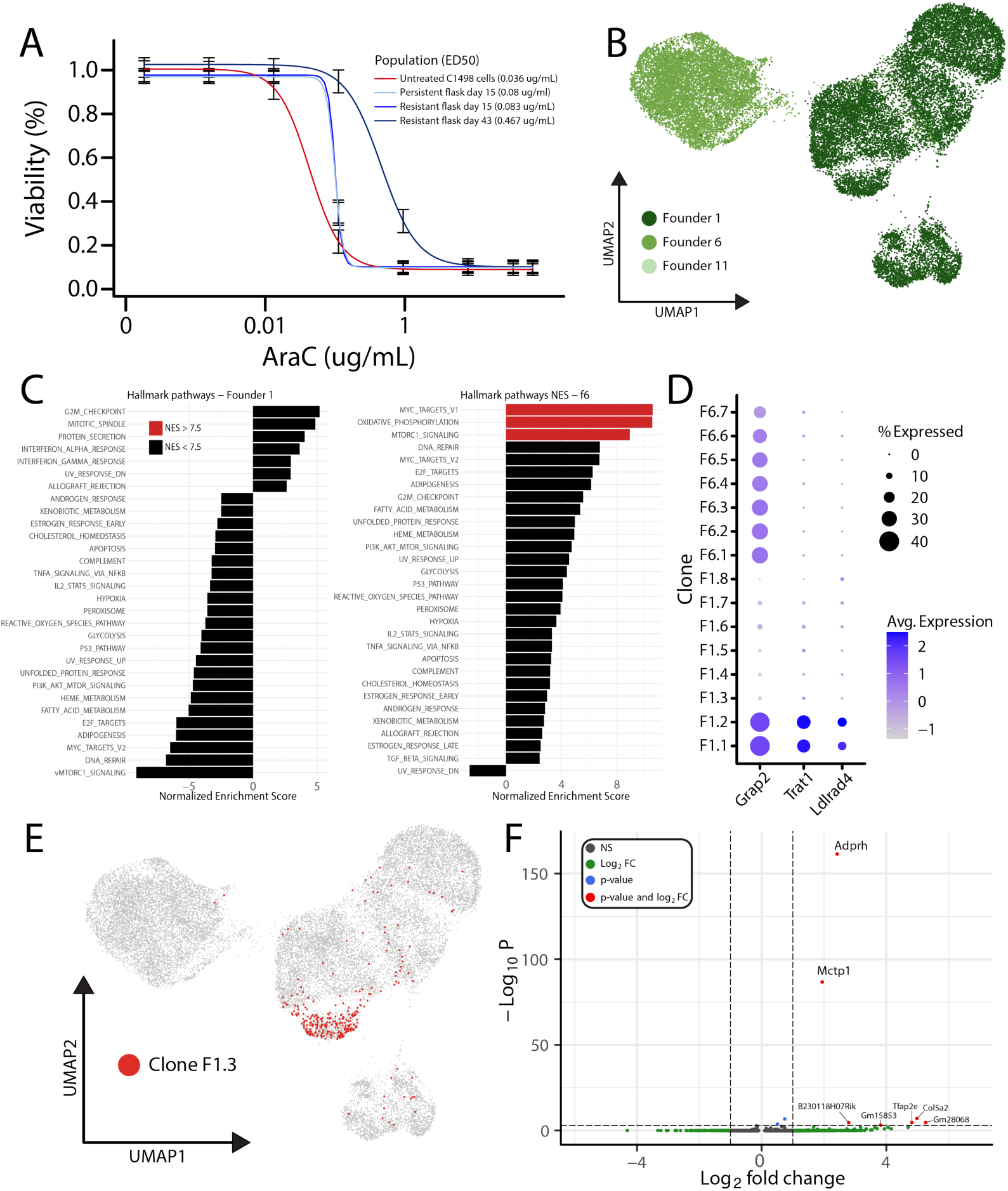
AraC treatment of C1498-FLARE cells in vitro leads to the generation of AraC-insensitive populations. FLARE recorder analysis reveals founder and clone-level transcriptomic differences. (**A**) AraC dose-response curves for C1498-FLARE cells through in vitro treatment. Cells in the resistant flask displayed an increase in ED50 values throughout treatment, indicating the emergence of an AraC-resistant population. Cells in the persistent flask also displayed an increase in ED50 values on day 15 of AraC treatment, but did not recover enough for another dose-response curve on day 43, suggesting higher AraC sensitivity compared to the resistant flask. (**B**) Captured C1498-FLARE cells colored by founder population. (**C**) GSEA analysis on founder 1 and founder 6 control cells to identify basal founder-specific differences. Analysis was conducted using the fGSEA R package and Hallmark gene sets from the mSigDB. (**D**) Dot plot showing expression of Grap2, Trat1, and Ldlrad4 in C1498-FLARE clone populations. (**E**) UMAP of C1498-FLARE cells with clone F1.3 cells highlighted in red. (**F**) Differential gene expression analysis on clone F1.3 cells compared to all founder 1 control cells uncovers clone-specific increased expression of Adprh and Mctp1.

**Supplemental figure 3:**
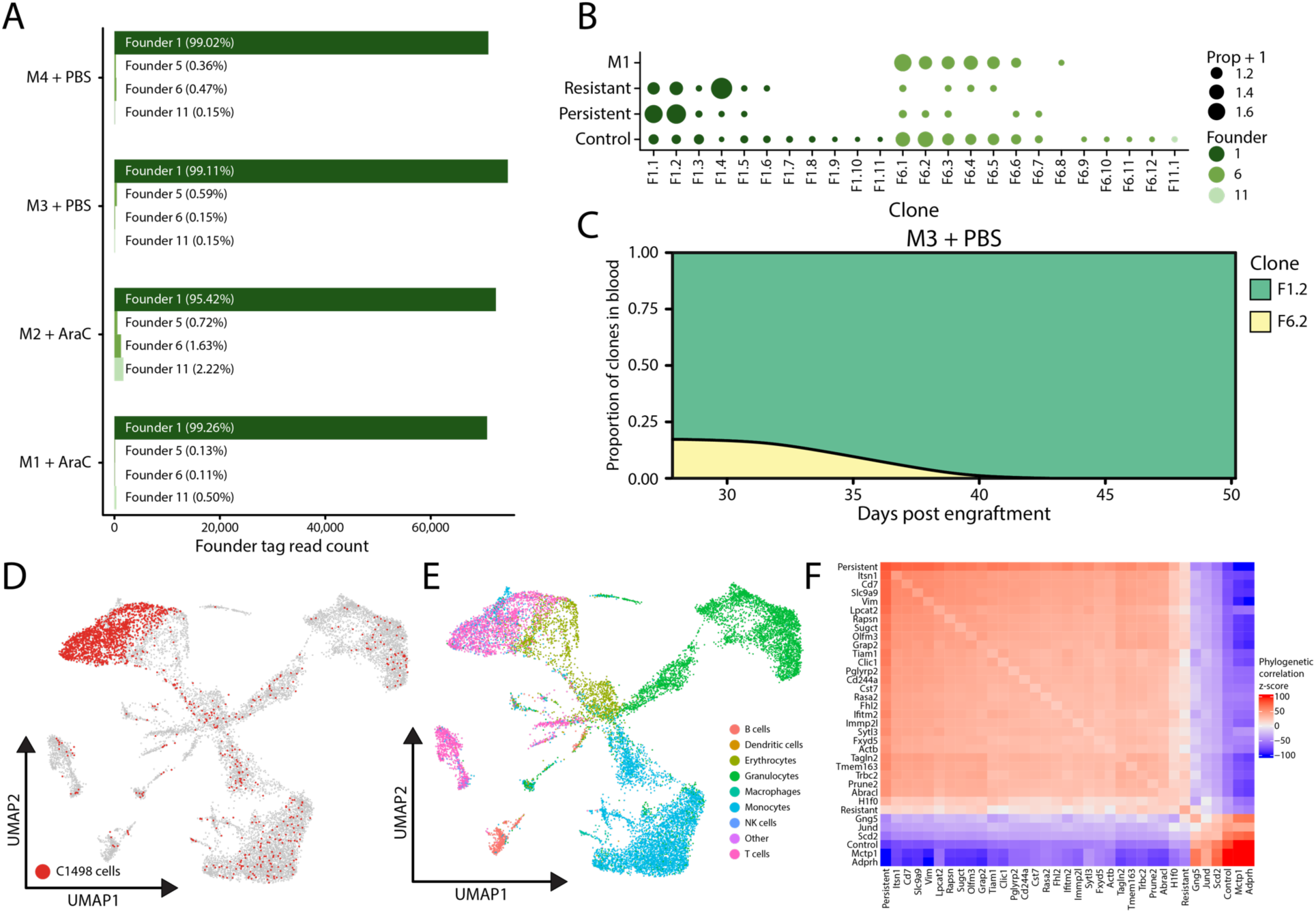
AraC treatment of C1498-FLARE cells in vivo leads to outgrowth of founder 6 cells. (**A**) Founder tag sequencing of blood draws taken 28 days after mice were injected IV with C1498-FLARE cells. (**B**) Clone populations for each sample, including in vivo sequencing from M1, first ordered by founder then by clone size. (**C**) Bulk sequencing of clone tags detected in blood draws from mouse M3. Clone tag reads were used to determine the clonal proportions of leukemic burden in the blood from day 30 to euthanasia on day 50. Founder proportions differ from figure 2E due to subsetting for clone tags detected in single cell data. (**D**) Captured single cells from endpoint blood and bone marrow collected from mouse M1. (**E**) Captured single cells from endpoint blood and bone marrow collected from mouse M1. Predicted C1498-FLARE cells were defined as having sufficient founder and clone tag reads to assign lineage identities, highlighted in red. (**F**) Cross-correlation heatmap of the top 35 auto-correlated genes over the Founder 1 tree.

**Supplemental figure 4:**
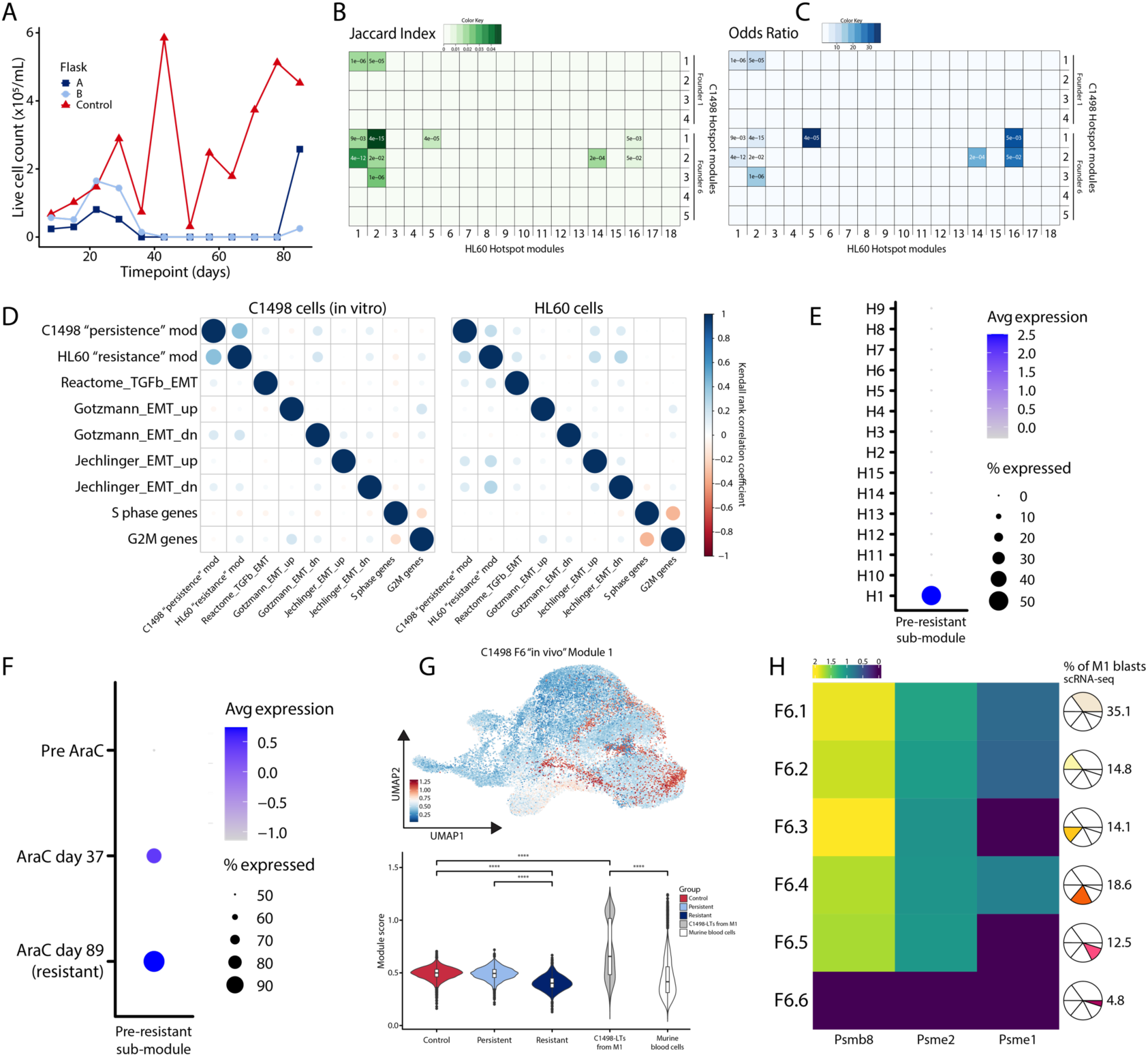
AraC treatment of HL60-FLARE cells in vitro leads to the generation of chemoresistance. Resistance-associated heritable Hotspot modules in C1498-FLARE and HL60-FLARE cells suggest the existence of drug response pathways common to murine and human AML cells. (**A**) Live cell counts for HL60-FLARE cells undergoing in vitro AraC treatment. Each flask was counted in duplicates at each timepoint, and counts were averaged. Control flask counts vary due to culture splitting in the days prior to counting. Treated flasks A and B did not have enough cells to be detected between day 40 and 80 and were not split during that time frame. (**B**) Jaccard Index similarity comparing C1498-FLARE and HL60-FLARE Hotspot modules. Square color is the Jaccard Index; values indicate BH-corrected P-values. (**C**) The odds ratio of gene overlaps comparing C1498-FLARE and HL60-FLARE Hotspot modules. Square color determined by odds ratio; values indicate BH-corrected P-values. (**D**) Expression scores for C1498 “persistence” Founder 1 module 1, HL60 “resistance” module 2, various EMT gene sets, and cell cycle gene sets are minimally correlated as measured by Kendall ranked correlation coefficient in C1498-FLARE and HL60-FLARE cells in vitro. (**E**) Dot plot showing expression of the pre-resistant sub-module in pre-experiment HL60-FLARE cells by founder. Founder H1 shows distinct upregulation of the signature. (**F**) Dot plot showing expression of the pre-resistant sub-module in AraC-treated HL60-FLARE H1 cells pre-experiment, at midpoint, and at endpoint. Pre-resistant signature expression increases through AraC treatment. (**G**) UMAP of founder 6 module 1 expression in the in vitro C1498-FLARE cells integrated with M1 endpoint sequencing. Below is expression quantification by cell group and shows module expression enrichment in cells collected from M1. (**H**) Log2 fold change values of Psmb8, Psme1, Psme2 genes for C1498-FLARE founder 6 clones comparing in vitro to in vivo clone cells.

**Supplemental figure 5:**
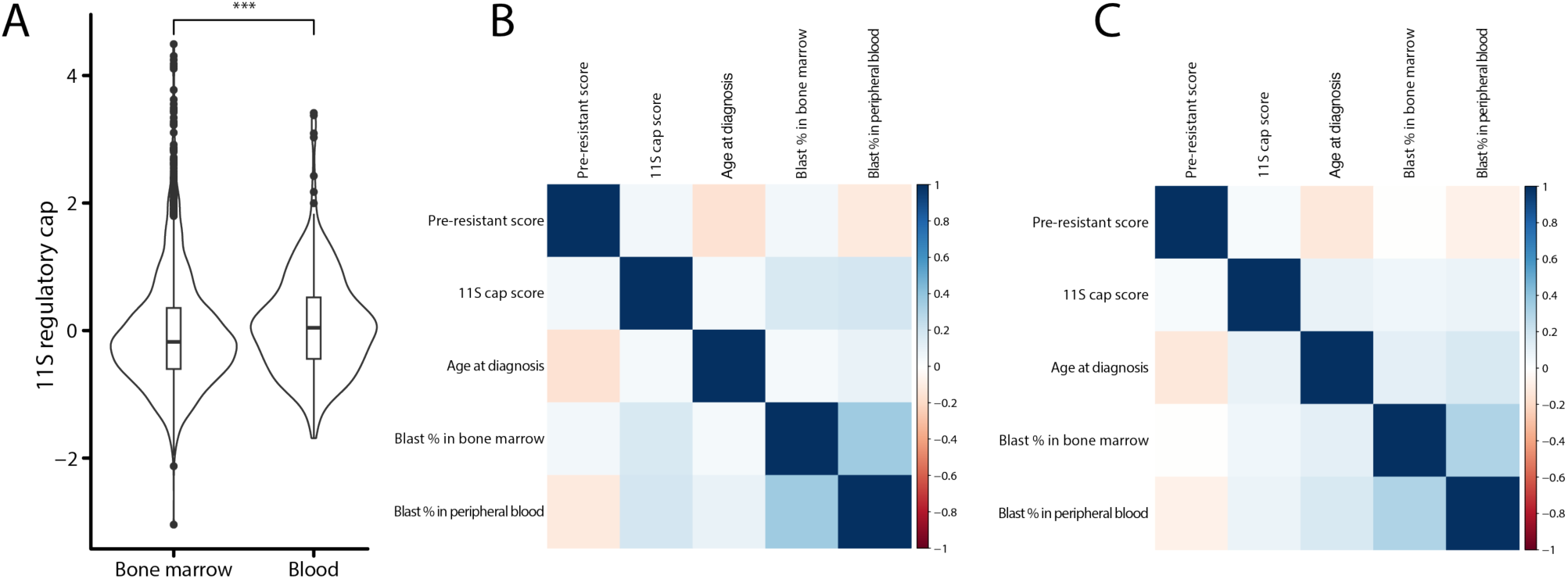
Expression of the 11S cap and pre-resistant signature are not strongly correlated with bone marrow or peripheral blood blast percentage at diagnosis. (**A**) Violin plot showing 11S regulatory cap gene set scores calculated from TARGET-AML patient samples taken at diagnosis. Mean expression in blood is significantly higher than in bone marrow. (**B**) TARGET-AML expression scores for 11S regulatory cap and pre-resistant signature in the bone marrow at diagnosis, age at diagnosis, and percentage of leukemic blasts in bone marrow and peripheral blood are minimally correlated as measured by Kendall ranked correlation coefficient. (**C**) TARGET-AML expression scores for 11S regulatory cap and pre-resistant signature in the peripheral blood at diagnosis, age at diagnosis, and percentage of leukemic blasts in bone marrow and peripheral blood are minimally correlated as measured by Kendall ranked correlation coefficient.

